# eRNAformer enables genome-wide *de novo* mapping of enhancer-derived RNA loci

**DOI:** 10.64898/2026.06.24.734403

**Authors:** Hua Yu, Wen Li, Wei Li, Yunjie Liu, Ying Chen, Xinwen Zhang, Siyue He, Zhiming Chen, Huiquan Wang, Jiajun Ni, Tingwei Gao, Fei Li, Lu Lu

## Abstract

Enhancer-derived RNAs (eRNAs) are critical regulators of gene transcription, yet their genome-wide annotation remains challenging. Here, we present eRNAformer, a multi-modal deep learning framework that integrates convolutional neural networks with transformers, specifically designed to capture long-range genetic features associated with bidirectional transcription. This approach enables *de novo* mapping of eRNA loci using DNA sequence and aggregated conventional RNA-seq data. When evaluated on ENCODE datasets, eRNAformer demonstrated high sensitivity and specificity in discriminating known eRNA loci from non-eRNA loci. Notably, the newly identified eRNA loci were enriched with evolutionarily constrained variants and genetic risk factors for complex diseases, and exhibit potential relevance for cancer therapy. Applied to GEO datasets, eRNAformer identified a range from 14,219 to 56,451 eRNA loci across multiple hematologic malignancies, facilitating the construction of a comprehensive eRNA database for blood cancers. We further identified and experimentally validated *FOXO1e*, a cluster of eRNAs located approximately 120 kb upstream of *FOXO1*, a known oncogene that drives t(8;21) acute myeloid leukemia (AML) preleukemic program. Together, these findings establish eRNAformer as a powerful tool for genome-wide eRNA annotation, provide a valuable resource for eRNA studies in hematologic cancers, and underscore the functional importance of eRNAs in AML pathogenesis.

## Introduction

Transcriptional regulation is orchestrated by the coordinated action of *trans*-acting factors, such as transcription factors, and *cis*-regulatory elements, including promoters and enhancers. While promoters act proximally to initiate transcription and generate stable RNA products, enhancers exert distal control over their target genes, often in a cell-type- and contex-specific manner. Active enhancers are typically marked by specific histone modifications, including H3K4 monomethylation (H3K4me1) and H3K27 acetylation (H3K27ac) [1]. Intriguingly, enhancers themselves are pervasively transcribed, giving rise to a class of non- coding RNAs known as enhancer- derived RNAs (eRNAs) [2–4]. Unlike long non-coding RNAs, eRNAs are predominantly unspliced, bidirectionally transcribed, and short-lived, yet they play critical roles in facilitating enhancer–promoter looping, chromatin remodeling, and the activation of target genes [5, 6]. Over the past decade, large- scale initiatives have systematically catalogued enhancer expression across thousands of cancer specimens and diverse human tissues, revealing the widespread involvement of eRNAs in development, homeostasis, and disease [7–9].

However, a fundamental and often overlooked limitation underlies virtually all these landmark studies: they uniformly rely on pre- existing enhancer annotations defined by histone modification marks such as H3K4me1 or H3K27ac. Consequently, eRNA quantification is performed directly on these pre- annotated intervals, which cannot effectively eliminate transcriptional noise and non-eRNA transcriptional signals originating from spurious RNA polymerase II occupancy at chromatin-accessible but non-functional regions. This issue is particularly acute in cancer contexts, where the transcriptional landscape is highly permissive and promiscuous, generating abundant background signals that obscure true enhancer activity. As a result, nascent RNA- seq technologies—including GRO- seq, PRO- seq, and their variants—remain the only reliable gold standard for genuine eRNA identification, as they capture actively transcribing RNA polymerases at base- pair resolution and enable *de novo* detection with high precision [10–15]. Yet these methods are technically demanding and resource-intensive, requiring large amounts of starting material, metabolic labeling, and specialized library preparation, which makes large- scale cancer profiling largely infeasible [16].

Several computational efforts have attempted to address this bottleneck using epigenetic or conventional RNA- seq data. DeepITEH [17] employs a deep-learning architecture that integrates DNA sequence and histone modification profiles to identify tissue- specific eRNAs. Separately, we previously developed eRNAFinder [18], which exploits bidirectional transcription patterns evident in RNA- seq read coverage to distinguish eRNAs from background. Although both methods represent important advances, they share a critical constraint: they operate exclusively on pre- annotated enhancer intervals. eRNAFinder requires predefined enhancer coordinates to compute its bidirectional signal statistics; DeepITEH relies on epigenomic features that are themselves derived from known enhancer marks. Consequently, neither approach can discover eRNAs originating from unannotated, novel, or context- specific enhancer loci. This limitation is particularly serious in cancer, where enhancer reprogramming frequently generates previously uncharacterized regulatory elements, and it underscores the pressing need for a method that can unlock the full potential of routine RNA- seq data for unbiased eRNA discovery.

Here we present eRNAformer, a multimodal deep-learning framework designed to perform *de novo* mapping of eRNA loci using DNA sequence and aggregated conventional RNA- seq data, entirely without any prior enhancer annotation. Rather than scoring predefined intervals, eRNAformer directly assembles transcripts from aggregated RNA- seq samples—merging reads across hundreds of datasets to generate a high- depth coverage profile. This aggregation- first strategy confers three key advantages: it consolidates recurrent bidirectional transcription signals while diluting stochastic noise, compensates for the extremely low abundance of eRNAs in individual samples, and operates without any external enhancer catalog. We curated 1955 RNA-seq samples from ENCODE across a wide range of tissues and cell types for model training (**Supplementary Dataset 1**). eRNAformer demonstrates superior sensitivity and specificity in distinguishing true eRNA loci from potential false positives, as rigorously validated through multiple tests using real eRNA loci data from the FANTOM5 and eRNAbase. The novel eRNAs mapped by eRNAformer are evolutionarily constrained and associated with complex diseases. Subsequent analysis revealed that the target genes of these novel eRNAs are significantly enriched in key oncogenic pathways and related to drug response and cancer therapy.

Applied to four substantial cohorts of over 2600 routine RNA-seq samples derived from acute myeloid leukemia (AML), chronic myeloid leukemia (CML), lymphoma (LM) and multiple myeloma (MM), eRNAformer successfully identified a range from 14219 to 56451 novel eRNA loci, resulting in the development of a blood-tumor-specific eRNA database. Furthermore, we experimentally validated the role of a cluster of eRNAs in regulating AML oncogenesis by activating *FOXO1*, confirming its biological relevance using both CRISPR inhibition and overexpression assays.

## Results

### Overview of eRNAformer

A defining characteristic of discovered eRNA loci is their bidirectional transcription, a pattern fundamentally determined by distinctive DNA sequences [19]. Based on this rationale, we designed the eRNAformer methodology to comprehensively capture the intrinsic DNA sequence features and corresponding bidirectional transcription patterns that accurately distinguish true eRNA loci from non-eRNA loci. eRNAformer integrates base-resolution DNA sequences with aggregated conventional RNA-seq signals as inputs to decode bidirectional transcription and elucidate the sequence-based genetic rules governing this divergent initiation process. To learn intricate feature dependencies both within and across these data modalities, we adopted the transformer architecture—widely used in natural language processing for its attention mechanisms that capture long-range dependencies—as a main partition of our modeling framework. To enable *de novo* mapping of eRNA loci, eRNAformer was trained on datasets assembled from aggregated conventional RNA-seq reads, comprising true eRNA loci and non-eRNA loci (**Methods**).

eRNAformer processes genomic inputs sequentially through a multi-scale convolutional block, a downsampling layer, a positional encoding module, a transformer tower, an attention pooling mechanism, and a binary classification head for eRNA locus prediction (**Fig. 1a, Extended Data Fig. 1a** and **Methods**). Specifically, multi-channel genomic inputs are first processed using parallel multi-scale 1-Dimension (1D) convolutions with varying kernel sizes. The resulting features are concatenated, then passed through batch normalization, Gaussian Error Linear Unit (GELU) activation, and dropout. This feature-extracted sequence undergoes length reduction via strided convolution before receiving relative positional encoding. A transformer encoder then models global dependencies between genetic elements of bidirectional transcription. The transformer outputs are adaptively compressed into a fixed-length representation through a learned attention pooling mechanism. This mechanism computes position-specific weights using tanh-activated linear layers followed by softmax normalization. Finally, a classification head comprising two linear layers (with Rectified Linear Unit (ReLU) activation and dropout applied between them) generates the prediction, where a sigmoid activation produces the prediction probability. This hybrid design effectively combines local feature extraction with global context modeling for accurate discovery of true eRNA loci.

**Fig. 1.**
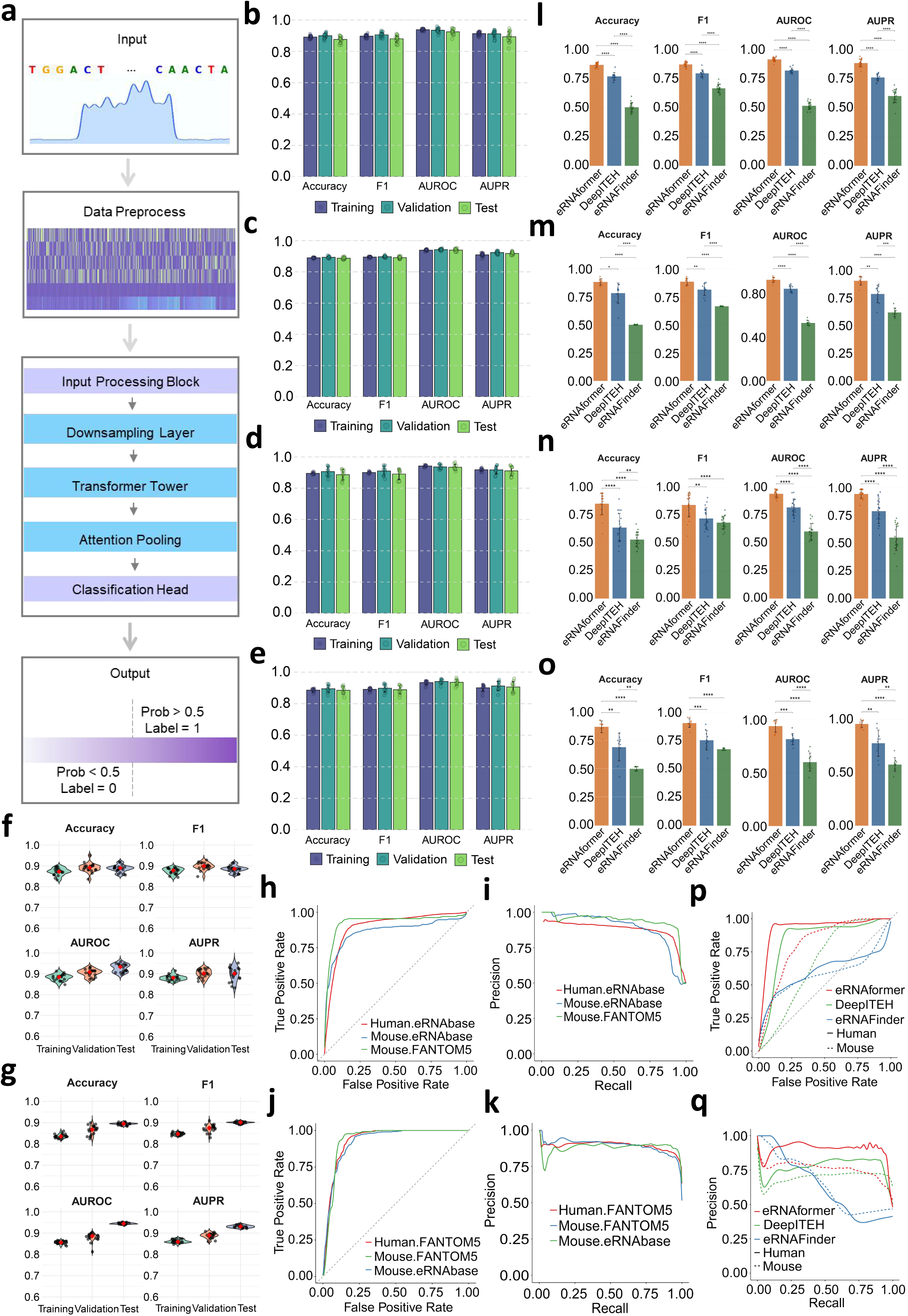
Model design and performance of eRNAformer. **a)** Overview of the eRNAformer model. **b)** Prediction performance of the eRNAformer model constructed on the human FANTOM5-derived benchmark dataset across 20 random data splits. **c)** Prediction performance of the eRNAformer model constructed on the human eRNAbase-derived benchmark dataset across 20 random data splits. **d)** Prediction performance of the eRNAformer model constructed on the human FANTOM5-derived benchmark dataset in 10-fold cross-validation experiments. **e)** Prediction performance of the eRNAformer model constructed on the human eRNAbase-derived benchmark dataset in 10-fold cross-validation experiments. **f)** Prediction performance of the eRNAformer model constructed on the human FANTOM5-derived benchmark dataset across 20 independently generated non-eRNA loci sets. **g)** Prediction performance of the eRNAformer model constructed on the human eRNAbase-derived benchmark dataset across 20 independently generated non-eRNA loci sets. **h)** ROC curves of the eRNAformer model constructed on the human FANTOM5-derived benchmark dataset for intra- and inter-species external data predictions. **i)** PR curves of the eRNAformer model constructed on the human FANTOM5-derived benchmark dataset for intra- and inter-species external data predictions. **j)** ROC curves of the eRNAformer model constructed on the human eRNAbase-derived benchmark dataset for intra- and inter-species external data predictions. **k)** PR curves of the eRNAformer model constructed on the human eRNAbase-derived benchmark dataset for intra- and inter-species external data predictions. **l)** Comparison of prediction performance between eRNAformer and other methods on the human FANTOM5-derived benchmark dataset across 20 random data splits; *p*-value < 0.0001(****), two-sided Wilcoxon signed-rank test. Data are presented as mean ± standard error of the mean (SEM). **m)** Comparison of prediction performance between eRNAformer and other methods on the human FANTOM5-derived benchmark dataset in 10-fold cross-validation experiments; *P*-value < 0.0001(****), *p*-value < 0.001(***), *p*-value < 0.01(**), *p*-value < 0.05(*), two-sided Wilcoxon signed-rank test. Data are presented as mean ± SEM. **n)** Comparison of prediction performance between eRNAformer and other methods on the mouse FANTOM5-derived benchmark dataset across 20 random data splits; *p*-value < 0.0001(****), *p*-value < 0.01(**), two-sided Wilcoxon signed-rank test. Data are presented as mean ± SEM. **o)** Comparison of prediction performance between eRNAformer and other methods on the mouse FANTOM5-derived benchmark dataset in 10-fold cross-validation experiments; *p*-value < 0.0001(****), *p*-value < 0.001(***), *p*-value < 0.01(**), two-sided Wilcoxon signed-rank test. Data are presented as mean ± SEM. **p)** ROC curves of eRNAformer and other methods on the human and mouse FANTOM5-derived benchmark dataset in 10-fold cross-validation experiments. **q)** PR curves of eRNAformer and other methods on the human and mouse FANTOM5-derived benchmark dataset in 10-fold cross-validation experiments.

### eRNAformer exhibits superior performance in identifying *bona fide* eRNA loci

Using known eRNA annotations from FANTOM5 [20] and eRNAbase [21] as references, we compiled two benchmark datasets comprising true eRNA loci and non-eRNA loci in the human and mouse genomes (**Methods** and **Supplementary Dataset 2**). These loci are comprehensively covered by aggregated conventional RNA-seq reads. On these benchmark datasets, eRNAformer demonstrates excellent performance in distinguishing eRNA loci from non-eRNA loci. This is evidenced by Areas Under Receiver Operating Characteristic (AUROC) and Precision-Recall (AUPR) curves in 20 random data splits and 10-fold cross-validation experiments (**Fig. 1b-e, Extended Data Fig. 1b-e**). The predictive capability of eRNAformer was further supported by high F1 score and accuracy (**Fig. 1b-e, Extended Data Fig. 1b-e**). Notably, the performance of eRNAformer models remains robust across different human and mouse benchmarking datasets (**Fig. 1b-e, Extended Data Fig. 1b-e**). To account for the random selection of non-eRNA loci regions inherent in the benchmark construction, we performed an additional evaluation using 20 independently generated non-eRNA loci sets. Consistently, eRNAformer maintains high performance for distinguishing true eRNA loci from non-eRNA loci (**Fig. 1f, 1g, Extended Data Fig. 1f, 1g**).

To further validate the generalizability of eRNAformer, we performed multiple external validations using independent benchmarking datasets derived from different databases and species. Specifically, models trained on one benchmark dataset were used to predict sample labels in another distinct benchmark dataset. As shown in **Fig 1h**-**k** and **Extended Data Fig. 1h**-**k**, eRNAformer accurately predicts the true labels across these datasets with high sensitivity and specificity. These results collectively indicate the conservation of bidirectional transcription rules between human and mouse, fully demonstrating the strong generalizability of the eRNAformer model in eRNA locus prediction. The consistently suboptimal performance of models in predicting the true labels of the eRNAbase-derived benchmarking dataset (**Fig 1h**-**k****, Extended Data Fig. 1h**-**k**) indicates the likely presence of false-positive eRNA locus annotations within the eRNAbase database. To comprehensively benchmark eRNAformer’s performance, we further compared its predictions with two existing methods, eRNAFinder [18] and DeepITEH [17], using FANTOM5-derived benchmark dataset. As measured by accuracy, F1 score and AUC, eRNAformer achieved the highest performance across all metrics and demonstrated statistically significant improvements over all competing methods in 20 random data splits and 10-fold cross-validation experiments (**Fig 1l-q**). The superior performance of models trained on the human dataset, compared to those trained on the mouse dataset (**Fig 1h-k**, **Fig 1p, q, Extended Data Fig. 1h-k**), indicates that increasing the volume of training data can lead to better predictive performance and generalizability. Taken together, our comprehensive validation and benchmarking comparison unequivocally show that eRNAformer achieves state-of-the-art performance and provides a reliable computational tool for accurately identifying eRNA loci across datasets and species.

### Combining DNA sequence and RNA-seq data synergistically enhances the prediction of eRNA loci

We next sought to determine the individual and combined contributions of DNA sequence and RNA-seq signals to the predictive performance of eRNAformer. To ensure the validity of the results, we utilized the model trained from FANTOM5 benchmark dataset to perform this analysis. Beyond the full multi-input eRNAformer architecture—which simultaneously incorporates both DNA sequence and RNA-seq data—we also trained and evaluated two ablated variants: one using only sequence information and another relying solely on RNA-seq signals. The full model consistently outperformed both single-input alternatives, achieving significantly higher predictive performance in validation and test datasets (**Fig 2a, 2c**). This trend was consistently observed across both the human and mouse genomes and was further supported by the binary cross-entropy (BCE) loss (**Fig 2a-d** and **Extended Data Fig 2a-f**). These results indicated that eRNAformer leverages the synergistic interplay between sequence context and RNA-seq signals to inform its decision-making process, enabling it to accurately discriminate functional eRNA loci from non-eRNA sequences. It is also notable that the performance gap between the full model and the two single-input models widened with increasing training sample size from mouse dataset (n=266) to human dataset (n=1,706) (**Fig 2a-d**, **Extended Data Fig 2a-d**). Moreover, these patterns were once again observed in external independent validation experiments (**Fig 2e-f**, **Extended Data Fig 2g-h**), highlighting that integrating DNA sequence and RNA-seq signal can capture more intrinsic features characterizing bidirectional transcription than single-input alternatives when training an eRNAformer model on large dataset.

**Fig. 2.**
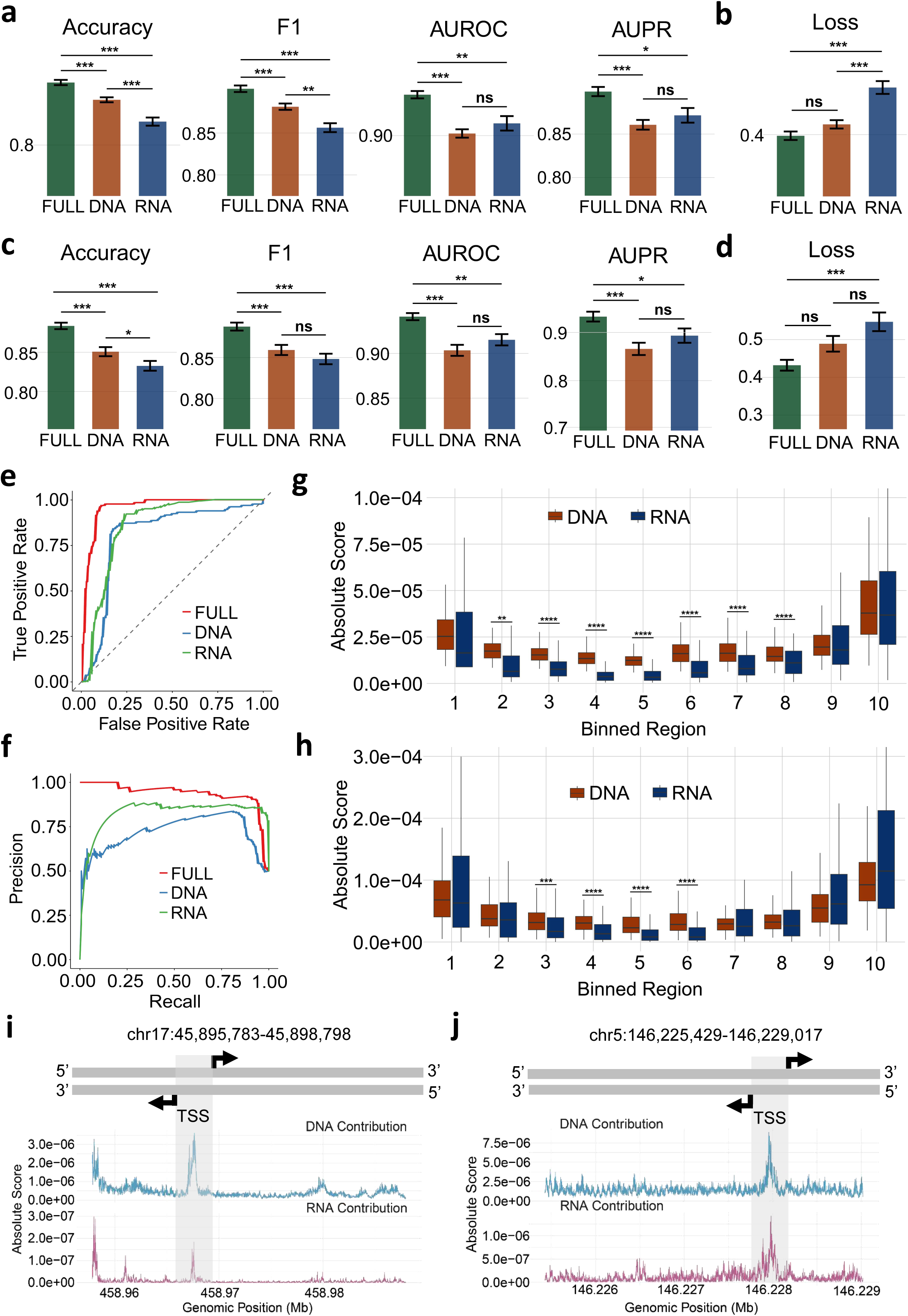
Combining DNA sequence and RNA-seq data synergistically improves the prediction of eRNA loci. **a)** Performance comparison between the full model and single-input models on the human validation set across 20 random data splits; *p*-value < 0.001 (***), *p*-value < 0.01 (**), *p*-value < 0.05 (*), *p*-value ≥ 0.05 (ns), two-sided Wilcoxon signed-rank test. Data are presented as mean ± SEM. **b)** BCE loss comparison between the full model and single-input models on the human validation set across 20 random data splits; *p*-value < 0.001 (***), *p*-value ≥ 0.05 (ns), two-sided Wilcoxon signed-rank test. Data are presented as mean ± SEM. **c)** Performance comparison between the full model and single-input models on the human test set across 20 random data splits; *p*-value < 0.001 (***), *p*-value < 0.01 (**), *p*-value < 0.05 (*), *p*-value ≥ 0.05 (ns), two-sided Wilcoxon signed-rank test. Data are presented as mean ± SEM. **d)** BCE loss comparison between the full model and single-input models on the human test set across 20 random data splits; *p*-value < 0.001 (***), *p*-value ≥ 0.05 (ns), two-sided Wilcoxon signed-rank test. Data are presented as mean ± SEM. **e)** ROC curves of the full model and single-input models, trained on the human FANTOM5-derived benchmark dataset, when predicting on the mouse FANTOM5-derived benchmark dataset. **f)** PR curves of the full model and single-input models, trained on the human FANTOM5-derived benchmark dataset, when predicting on the mouse FANTOM5-derived benchmark dataset. **g)** Distribution of DNA and RNA contribution scores across 10 equally divided regions (5′ to 3′) in the human FANTOM5-derived benchmark dataset; *p*-value < 0.0001 (****), *p*-value < 0.01 (**), two-sided Wilcoxon signed-rank test. **h)** Distribution of DNA and RNA contribution scores across 10 equally divided regions (5′ to 3′) in the mouse FANTOM5-derived benchmark dataset; *p*-value < 0.001 (****), *p*-value < 0.001 (***), two-sided Wilcoxon signed-rank test. **i)** DNA and RNA contribution score profile along a representative human benchmark eRNA locus. **j)** DNA and RNA contribution score profile along a representative mouse benchmark eRNA locus.

To further investigate the contributions of the two input modalities in identifying genuine eRNA loci, we employed the integrated gradients method to quantify the contribution scores of the gene sequence and RNA-seq data, respectively (**Methods**). First, we evaluated the overall contributions of the gene sequence and RNA-seq signals across the benchmark eRNA regions. By analyzing 20 eRNAformer models trained on independently generated sets of non-eRNA loci, we found that the DNA sequence modality contributed significantly more to eRNA locus prediction than the RNA-seq signal modality (**Extended Data Fig. 2i**). This observation aligns with the biological fact that bidirectional transcription is primarily determined by the genetic nucleic acid sequence. To gain a deep understanding of sequence-mediated regulation on bidirectional transcription initiation and termination, we next examined the positional dependency of DNA and RNA contributions (**Methods**). The analysis revealed that both DNA and RNA exhibited the highest contribution scores at the start and end positions of the eRNA regions (**Extended Data Fig 2j, k**), indicating that eRNAformer successfully captured the termination signals of bidirectional transcription.

Notably, relatively high contribution values from both DNA and RNA were also detected at the center of the eRNA regions (**Extended Data Fig 2j, k**). This pattern was more pronounced in the relatively smaller mouse dataset (**Extended Data Fig2j, k**), suggesting that eRNAformer recognized the initiation signals of bidirectional transcription. Interestingly, DNA contributions in the central eRNA region were substantially greater than those of RNA (**Fig 2g, h**), implying that the key genetic codes for bidirectional transcription embedded in the DNA sequence are crucial for eRNAformer to capture complex dependencies within and across modalities.

To intuitively demonstrate the value of integrating DNA and RNA information for capturing sequence elements that determine bidirectional transcription and ensure precise eRNA locus prediction, we examined two benchmark eRNA loci that were successfully identified by the full eRNAformer model but missed by single-input alternatives. First, we analyzed contribution scores along a human benchmark eRNA locus (chr17:45,895,783–45,898,798). The results show that the contribution profiles of the DNA sequence align closely with those of the RNA-seq signal (**Fig 2i**). Importantly, distinct peaks in both DNA and RNA contributions were observed at the initiation region of bidirectional transcription, in contrast to the surrounding regions (**Fig 2i**). We also investigated a mouse benchmark eRNA locus (chr5:146,225,429–146,229,017) and found that regulatory sequences near the transcription initiation region were robustly captured, with clear co-enrichment of both modalities in this functional segment (**Fig 2j**). Collectively, these findings underscore the importance of integrating DNA sequence and RNA-seq data to accurately predict eRNA loci and decipher the regulatory logic of bidirectional transcription.

### eRNAformer maps evolutionarily constrained eRNAs associated with complex diseases

Having demonstrated the ability of eRNAformer in identifying *bona fide* eRNAs, we next employed eRNAformer to identify novel eRNAs from ENCODE aggregated RNA-seq samples. Given that incorporating more training data effectively enhances model predictive performance, we fine-tuned the model—originally trained on the human FANTOM5-derived benchmark dataset—by integrating the mouse FANTOM5-derived benchmark dataset.

Leveraging this fine-tuned model, we then identified potential eRNA loci in both the human and mouse genomes using aggregated reads from ENCODE RNA-seq samples. Of 14,661 human and 8,484 mouse assembled single-exon transcription loci, eRNAformer identified 8,809 and 2,812 as high-confidence eRNA loci (**Supplementary Dataset 3**). These putative eRNA regions and their flanking sequences displayed marked enrichments in H3K4me1 and H3K27ac signals compared to non-eRNA loci across both genomes (**Extended Data Fig 3a-d**), suggesting their potential biological functionality. Since functional eRNA loci are often under evolutionary constraint, we next evaluated the evolutionary conservation of identified eRNA loci on human genome by analyzing both intra-species genetic variation and cross-species constraint. To assess intra-species variation, we mapped variants from the Genome Aggregation Database (gnomAD) [22]—the largest catalog of human genetic variation—to the identified eRNA loci and their flanking regions of equal length. The number of variants falling within these three genomic compartments was systematically quantified. We observed that eRNA loci consistently harbored significantly fewer variants compared to their flanking regions across multiple chromosomes. This pattern remained highly robust when stratifying variants by minor allele frequency (MAF) (**Fig 3a, Extended Data Fig 3e**). We subsequently examined interspecies evolutionary constraint by calculating PhyloP scores for variants within these regions. Consistently elevated PhyloP scores were observed across all MAF strata (**Fig 3b, Extended Data Fig 3f**), indicating stronger evolutionary conservation. Together, these results demonstrate that eRNAformer identifies functional eRNAs under clear evolutionary constraints.

**Fig. 3.**
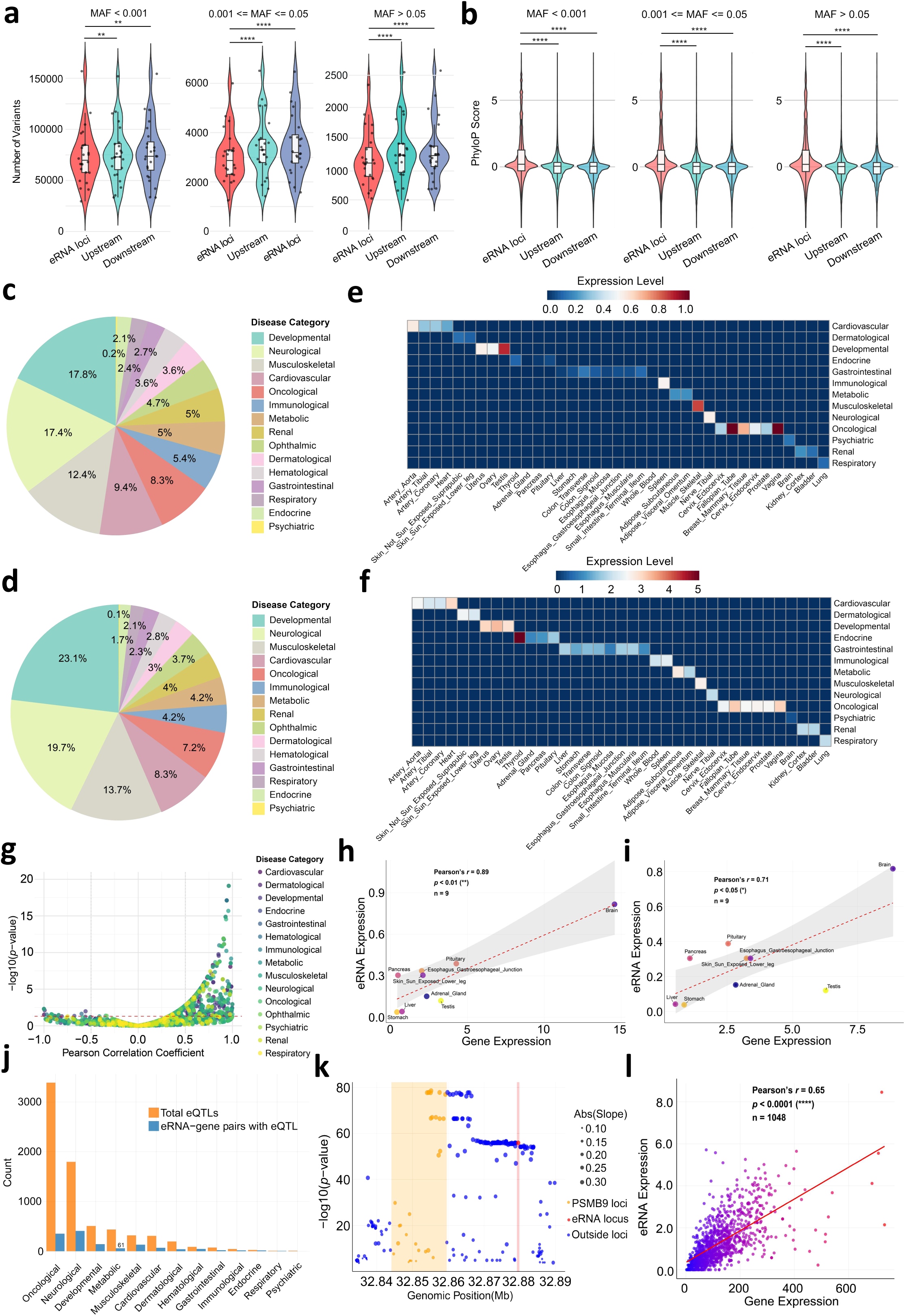
eRNAformer maps evolutionarily constrained eRNAs associated with complex diseases. **a)** Violin plots with boxplots displaying the distribution of variant counts in eRNA loci predicted by eRNAformer and in flanking regions of equal length. Variants are stratified into three groups based on minor allele frequency (MAF), with each point representing the number of variants on a single chromosome; *p*-value < 0.0001 (****), *p*-value < 0.01 (**), two-sided Wilcoxon signed-rank test. **b)** Violin plots with boxplots illustrating the distribution of PhyloP conservation scores for variants located within eRNAformer-predicted eRNA loci and comparable flanking regions. Variants are categorized into three groups according to MAF; *p*-value < 0.0001 (****), two-sided Wilcoxon signed-rank test. **c)** Pie chart showing the distribution of eRNAs associated with 15 distinct types of diseases. **d)** Pie chart depicting the distribution of target genes linked to 15 different disease categories. **e)** Heatmap representing the expression profiles of eRNAs across various tissues, specifically for eRNA-target gene pairs associated with a unique disease type. **f)** Heatmap depicting the expression patterns of target genes across multiple tissues, restricted to eRNA-target gene pairs linked to a single disease category. **g)** Scatter plot displaying the Pearson correlation coefficient versus the -log(*p*-value) for the expression of eRNA-target gene pairs. Points are colored according to the disease type associated with each pair. The red horizontal line indicates the significance threshold. **h)** Scatter plot showing the correlation of expression across different tissues between the eRNAformer-identified eRNA (chr17:46,243,606-46,245,044) and its target gene *MAPT*. The symbol *n* denotes the number of tissue samples used in the correlation analysis. **i)** Scatter plot illustrating the cross-tissue expression correlation between the eRNAformer-predicted eRNA (chr17:45,975,492-45,980,506) and its target gene *KANSL1*. The symbol *n* denotes the number of tissue samples used in the correlation analysis. **j)** Bar plot showing the total number of eQTLs for target genes associated with different diseases, along with the number of these eQTLs located within the corresponding eRNA loci. Only eRNA-target pairs with significant correlations are included. **k)** Scatter plot demonstrating an eQTL of the target gene *PSMB9* located within the eRNA region (chr6:32,879,171-32,879,848). The area shaded in red denotes the eRNA locus, while the area in yellow corresponds to the gene region. **l)** Scatter plot illustrating the expression correlation between the eRNA (chr6:32,879,171-32,879,848) and its target gene PSMB9 in whole blood tissue.

Given the frequent association of functional eRNAs with diseases, we evaluated the potential disease relevance of these eRNA loci. We identified their putative target genes located within 1 Mb and interrogated the disease associations of these genes using the ClinVar database [23]. Our analysis revealed that 3192 target genes of 1635 eRNA loci are linked to 15 disease categories, with developmental disorders representing the most frequently associated category for both eRNAs and their target genes (**Fig 3c, 3d, Extended Data Fig 3g**). Notably, these eRNA loci exhibited overwhelmingly significant enrichment of H3K4me1 and H3K27ac marks compared to non-eRNA loci (**Extended Data Fig 3h, 3i**). Interestingly, eRNAs and their target genes linked to a single disease category showed striking tissue specificity, with their expression highly enriched in the corresponding disease-relevant tissues (**Fig 3e, 3f**), consistent with the established role of eRNAs in activating the transcription of their target genes. We investigated the correlation between eRNA expression and the expression of their target genes using transcriptomic data from GTEx dataset [24]. The results demonstrated that the vast majority of eRNAs exhibit highly significant positive expression correlation with their target genes, with only a limited number of negative correlations observed (**Fig 3g**). For instance, one eRNAs (chr17:45,969,467-45,969,967) showed significant positive correlation with the expression of its target genes *MAPT* and *KANSL1* (**Fig3h, 3i**). Both eRNAs and their target genes are highly expressed in brain tissues (**Fig3h, 3i**). Notably, *MAPT1* and *KANSL* have been previously implicated in neurological diseases [23].

To investigate the regulatory potential of the significantly correlated eRNA-target gene pairs, we analyzed whether the expression quantitative trait loci (eQTLs) of the target genes were located within their corresponding eRNA loci. Our analysis revealed 1,326 eRNA-target pairs where the eQTLs for the target genes were colocalized with the eRNA loci (**Fig 3j**). For example, the eQTLs for the immune-related genes *PSMB9* and *CCL5*, as well as for a gene SYK associated with skin disease, were located within their corresponding eRNA loci, and these pairs showed significant expression correlation in disease-relevant tissues (**Fig 3k, 3l, Extended Data Fig 3j-m**). Collectively, these results demonstrate that eRNAformer has successfully identified a substantial number of functionally important and disease-associated eRNA loci.

### eRNAformer discovers novel eRNAs that holds potential value for cancer therapy

Having established a link between eRNAformer-identified eRNAs and cancer (**Fig. 3c**), we next systematically analyzed their expression across multiple cancer types using publicly available TCGA data. To map the expression landscape of these novel eRNAs, we calculated RNA-seq signals from TCGA over eRNA regions and defined eRNAs with an average expression ≥ 0.01 TPM as detectable per cancer type. This approach identified 671 detectable eRNAs across cancers (**Supplementary Dataset 4**), ranging from 144 in UVM to 564 in STAD (**Extended Data Fig. 4a**). In contrast to earlier reports [7], most eRNAs were expressed in more than five cancer types (**Extended Data Fig. 4b**). Expression similarity analysis revealed strong cancer-type specificity, with samples clustering by cancer type (**Fig. 4a**). Cancers with similar histology also grouped together—such as squamous cell carcinomas (BLCA, CESC, HNSC, LUSC), kidney cancers (KIRC, KIRP, KICH), and neuronal cancers (GBM, LGG, PCPG) (**Extended Data Fig. 4c**)—indicating that these eRNAs can characterize molecular features across cancers.

**Fig. 4.**
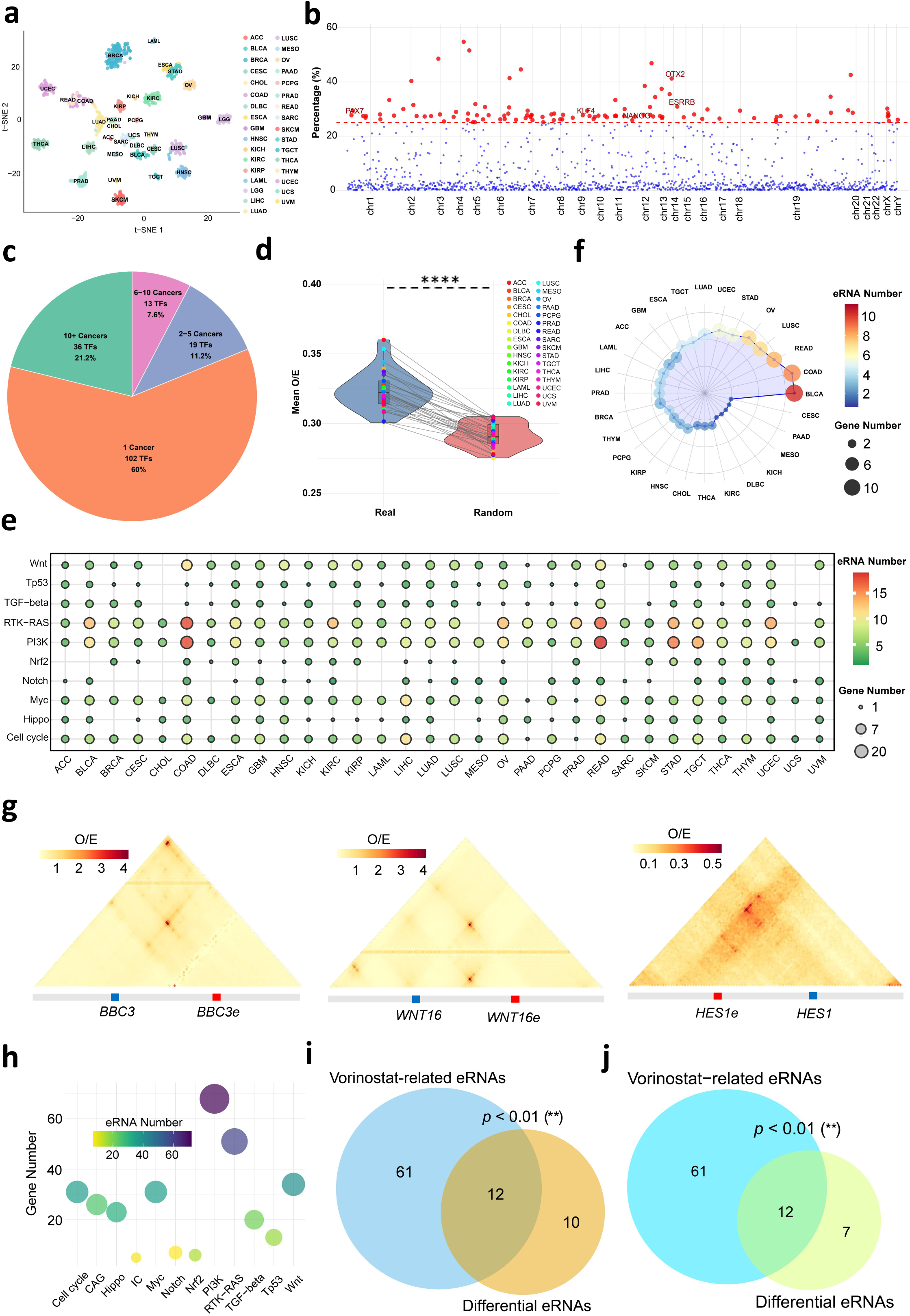
eRNAformer discovers novel eRNAs that holds potential value for cancer therapy. **a)** t-SNE plot illustrating cancer-type-specific patterns of novel eRNAs identified by eRNAformer. Dots represent cancer samples, and colors indicate cancer types. **b)** Scatter plot of master regulators of eRNAs in DLBC. Each dot corresponds to a transcription factor (TF). Red dots highlight master regulators significantly correlated with ≥25% of individual eRNAs. **c)** Pie chart depicting the distribution of master TFs based on the number of cancer types they regulate: 1 cancer type, 2–5 cancer types, 6–10 cancer types, and >10 cancer types. **d)** Violin plot comparing the averaged Hi-C observed/expected (O/E) contacts across cancer types between true eRNA–target links and distance-matched random pairs; *p*-value < 0.0001 (****), two-sided Wilcoxon signed-rank test. **e)** Droplet chart showing the number of eRNAs and target genes involved in Hi-C-supported regulatory links across different cancer types and signaling pathways. **f)** Radar chart displaying the number of eRNAs and clinically actionable genes (CAGs) included in Hi-C-supported regulatory links across various cancer types. **g)** Hi-C O/E contacts for three regulatory links between eRNAs and cancer signaling pathway genes. **h)** Droplet chart showing the number of genes significantly associated with CCLE anticancer drug sensitivity across different gene sets, along with the number of linked eRNAs. **i)** Venn diagram showing the overlap between eRNAs differentially expressed after Vorinostat treatment in melanoma cell lines and eRNAs targeting Vorinostat-associated genes. **j)** Venn diagram showing the overlap between eRNAs differentially expressed after Vorinostat treatment in melanoma patients and eRNAs targeting Vorinostat-associated genes.

We then identified putative eRNA regulators using co-expression and binding motif analysis between each eRNA and transcription factor (TF) (**Methods**). TFs correlated with ≥ 25% of eRNAs in a cancer were classified as master regulators. Using this method, we identified 170 master regulators across different cancer types (**Supplementary Dataset 4**). For example, 110 master regulators were identified in diffuse large B-cell lymphoma (DLBC) (**Fig. 4b, Extended Data Fig. 4d**), including known enhancer regulators such as PAX7, OTX2, KLF4, NANOG, and ESRRB [25–28]. Notably,55 of these master regulators were known cancer-related TFs in the Tfcancer database [29]. Most master regulators (102 of 170) correlated strongly with eRNAs in only one cancer type (**Fig. 4c**). We also found 36 master regulators that function in ≥ 10 cancer types; among these, 39% were linked to genomic instability (**Extended Data Fig. 4e**), consistent with reported associations between copy number alterations and enhancer activation [9].

We further constructed a pan-cancer global eRNA–gene regulatory network based on physical distance and co-expression (**Methods**). This analysis identified 611 putative target genes significantly correlated with 665 eRNAs in at least one cancer type. Using ENCODE Hi- C data, we examined three-dimensional chromatin interactions for all predicted eRNA–gene connections and found that more than 77% (18,575/24,124) were supported by significant Hi- C interactions (**Methods**). The proportion of Hi- C- supported connections was significantly higher than that of distance- matched random pairs across cancer types (**Fig. 4d, Extended Data Fig. 4f**). To investigate the regulatory roles of eRNAs in cancer, we compiled 1,056 cancer signaling pathway genes (CSPGs), 134 clinically actionable genes (CAGs), and 44 immune checkpoints (ICs). Among these, 198 genes were associated with 169 eRNAs in at least one cancer type, and 99% (1,395/1,404) of these eRNA–gene associations were supported by 3D chromatin interactions (**Fig 4e, f, Extended Data Fig 4g**) —for example, three in cancer signaling pathways, two in clinically actionable genes, and one in immune checkpoints (**Fig. 4g, Extended Data Fig. 4h, i**). By integrating gene expression and drug sensitivity (AUC) data from the Cancer Cell Line Encyclopedia (CCLE) and the Genomics of Drug Sensitivity in Cancer (GDSC), we further assessed correlations between the expression of Hi-C supported eRNA-linked genes and drug responses. The majority of these genes (190/198) showed significant correlations with anticancer drugs (FDR < 0.05; **Fig. 4h, Extended Data Fig. 4j**). Notably, eRNAs differentially expressed upon Vorinostat treatment in melanoma cell lines and patients [30] (**Methods**) were significantly enriched among eRNAs targeting Vorinostat- associated genes (**Fig. 4i, j**), implying a functional role for eRNAs in mediating drug response. Collectively, these findings demonstrate that novel eRNAs identified by eRNAformer are closely linked to cancer biology and anticancer drug sensitivity, highlighting their potential clinical relevance for cancer therapy.

### A multitude of cancer-related eRNAs are aberrantly activated in hematological malignancies

We next applied eRNAformer to *de novo* map eRNA loci in multiple hematological malignancies using aggregated RNA-seq reads. We collected 2665 RNA-seq samples of human and mouse hematological malignancies from the Gene Expression Omnibus (GEO) database (**Supplementary Dataset 5**). Using these samples, eRNAformer identified 146172 and 70358 novel eRNAs in hematological malignancies in human and mouse genomes, respectively (**Supplementary Dataset 5**). Notably, nearly all identified mono-exonic transcripts that overlapped with known eRNA loci in the FANTOM5 and eRNAbase databases were predicted to be authentic eRNAs (**Extended Data Fig 5a, b**). This result further confirms that eRNAformer has successfully learned the intrinsic features of eRNA loci, supporting its value for *de novo* eRNA locus mapping. Interestingly, the expression profiles of these novel eRNAs can effectively distinguish the different types of hematological malignancies and normal samples (**Fig.5a**, **Extended Data Fig 5c**). We next constructed regulatory networks that link these eRNAs to upstream TFs and downstream target genes across different hematological malignancies. The human hematological malignancy networks consist of 146096 eRNAs linked to 293 known cancer-associated TFs curated in Tfcancer database and 1104 known cancer-associated target genes (CSPGs, CAGs and ICs). Correspondingly, the mouse hematological malignancy networks consist of 70326 eRNAs linked to 266 known cancer-associated TFs and 1149 known cancer-associated target genes. Additionally, 98% and 95% of the eRNA-target gene regulatory connections in the human and mouse hematological malignancy networks, respectively, are supported by ENCODE Hi-C interactions.

To explore the functional relevance of these eRNAs across different hematologic malignancies, we first identified eRNAs that were consistently activated in all four types of hematologic tumors compared with normal samples. In the human genome, 898 eRNAs were highly expressed across all four tumor types (**Fig. 5b**). Accordingly, 488 target genes regulated by these eRNAs were also highly expressed in the same four tumor types (**Fig. 5b**). For instance, an eRNA (chr2:110,626,740-110,629,171) and the well- known cancer-related gene *BUB1* [31] showed significant expression correlations across all four hematologic tumors (**Fig. 5c**). Notably, a similar eRNA–target gene expression correlation pattern was also observed in mouse (**Extended Data Fig. 5c–e**). We next performed gene set variation analysis (GSVA) to compute GSVA scores for the eRNA- regulated target gene sets across different cancer samples. Based on these scores, samples were stratified into high, medium, and low groups. Ten hallmark gene sets associated with cancer development and progression [32] were then examined for differences in average expression between the high- score and low- score groups. Eight of the ten gene sets were significantly more highly expressed in the high- score tumor samples than in the low- score group, a pattern consistently observed in both human and mouse tumor samples (**Fig. 5d, e**). Consistent with these results, gene set enrichment analysis (GSEA) revealed marked expression differences in cancer signaling pathway genes between the high- and low- score groups. For example, cell cycle pathway genes were significantly upregulated in human high- score samples, whereas p53 signaling pathway genes were significantly downregulated in mouse high- score samples (**Extended Data Fig. 5f, g**). Interestingly, comparison of cell type abundances between the high- and low- score group samples showed that the high- score group had a significantly lower abundance of M1 macrophages (**Extended Data Fig. 5h, i**). As M1 macrophages drive pro-inflammatory responses and anti-tumor immunity, their reduced abundance suggests impaired immune activation and tumor cell clearance, likely contributing to a more immunosuppressive and tumor-permissive microenvironment. Together, these findings indicate that the consistently activated eRNAs play important functional roles in multiple hematologic malignancies.

**Fig. 5.**
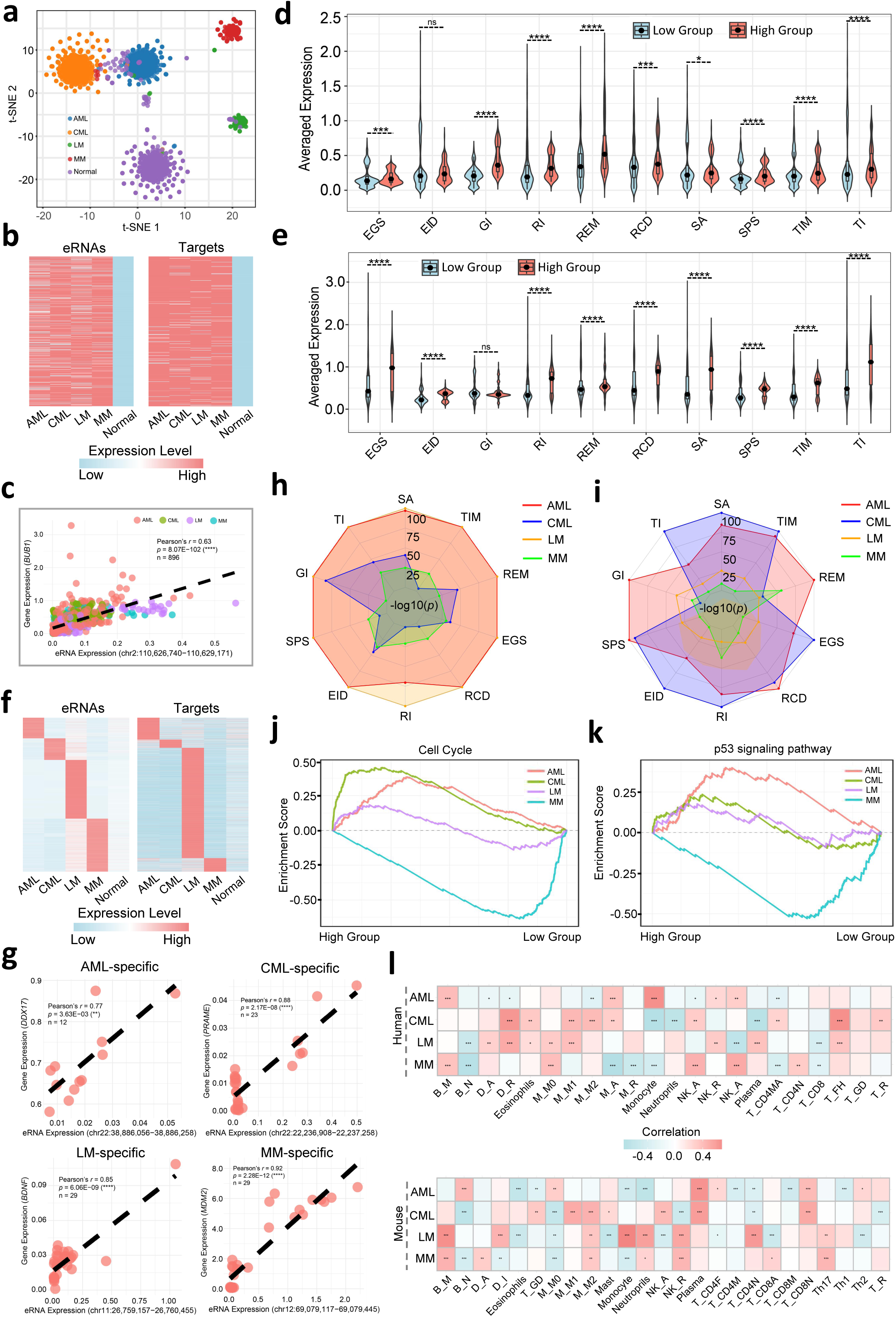
A multitude of cancer-related eRNAs are aberrantly activated in hematological malignancies. **a)** t-SNE plot showing the mapping of human hematologic tumor samples and normal samples onto the first two dimensions, with colors denoting sample types **b)** Heatmap displaying 898 consistently activated eRNAs and their 488 target genes across multiple human hematological malignancies. **c)** Scatter plot showing the expression correlation between a consistently activated eRNA (chr2:110,626,740-110,629,171) and the well-known cancer-related gene *BUB1* across different human hematological malignancies. The *p*-value was obtained using test of correlation coefficient. **d)** Violin plot showing the average expression differences of hallmark gene sets between the high-score and low-score groups, defined by the GSVA score of the target gene set regulated by consistently activated eRNAs in human hematological malignancies. GS: Growth Suppressors; EID: Evading Immune Destruction; GI: Genome Instability; RI: Replicative Immortality; REM: Reprogramming Energy Metabolism; RCD: Resisting Cell Death; SA: Sustained Angiogenesis; SPS: Sustaining Proliferative Signaling; TIM: Tissue Invasion and Metastasis; TI: Tumor-promoting Inflammation. *p*-value < 0.0001 (****), *p*-value < 0.001 (***), *p*-value < 0.05 (*), *p*-value ≥ 0.05 (ns), two-sided Wilcoxon signed-rank test. **e)** Violin plot showing the average expression differences of ten hallmark gene sets between the high-score and low-score groups, defined by the GSVA score of the target gene set regulated by consistently activated eRNAs in mouse hematological malignancies. *p*-value < 0.0001 (****), *p*-value ≥ 0.05 (ns), two-sided Wilcoxon signed-rank test. The *p*-values were obtained using test of correlation coefficient. **f)** Heatmap displaying 9,096 eRNAs and 4,800 target genes specifically expressed in distinct human hematological malignancies. **g)** Scatter plot showing the expression correlations between cancer-type-specifically expressed eRNAs and their target genes in distinct human hematological malignancies. The *p-*value was derived from a correlation coefficient test. **h)** Radar plot showing the distribution of *p*-values from differential expression analysis of ten hallmark gene sets between high and low groups stratified by cancer-specific expressed eRNA target genes in human hematological malignancies, using a two-sided Wilcoxon signed-rank test. **i)** Radar plot showing the distribution of *p*-values from differential expression analysis of ten hallmark gene sets between high and low groups stratified by cancer-specific expressed eRNA target genes in mouse hematological malignancies, using a two-sided Wilcoxon signed-rank test. **j)** GSEA plot displaying the expression changes of cell cycle genes between high and low groups stratified by cancer-specific expressed eRNA target genes in human hematological malignancies. **k)** GSEA plot displaying the expression changes of p53 signaling pathway genes between high and low groups stratified by cancer-specific expressed eRNA target genes in human hematological malignancies. **l)** Heatmap showing the correlation between GSVA scores of tumor-type-specific eRNA-regulated target gene sets and immune-related cell proportions across different cancer samples. The *p*-values were obtained using test of correlation coefficient. B_N, Naive B-cells; B_M, Memory B-cells; T_CD8, CD8 T-cells; T_CD4N, CD4 Naive T-cells; T_CD4MR, CD4 Memory Resting T-cells; T_CD4MA, CD4 Memory Activated T-cells; T_FH, Follicular Helper T-cells; T_R, Regulatory T-cells; T_GD, Gamma Delta T-cells; NK_R, Resting NK cells; NK_A, Activated NK cells; M_M0, M0 Macrophages; M_M1, M1 Macrophages; M_M2, M2 Macrophages; D_R, Resting Dendritic cells; D_A, Activated Dendritic cells; M_R, Resting Mast cells; M_A, Activated Mast cells; T_CD8A, CD8 Activated T-cells; T_CD8N, CD8 Naive T-cells; T_CD8M, CD8 Memory T-cells; T_CD4M, CD4 Memory T-cells; T_CD8N, CD8 Naive T-cells; T_CD4F, CD4 Follicular T-cells; D_I, Immature Dendritic cells.

To further investigate the tumor- specific functional relevance of these eRNAs in different hematologic malignancies, we next identified eRNAs that were exclusively activated in each individual tumor type. In the human genome, 1222, 1273, 3130 and 3471 eRNAs were specifically activated in AML, CML, LM, and MM, respectively (**Fig. 5f**). Accordingly, 696, 262, 3510 and 432 target genes regulated by these eRNAs were highly expressed in the corresponding tumor types (**Fig. 5f**). For example, significant expression correlations were observed for the following eRNA–target gene pairs: an eRNA (chr22:38,886,056-38,886,258) with the known AML- associated gene *DDX17* [33]; an eRNA (chr22:22,236,908-22,237,258) with the known CML- associated gene *PRAME* [34]; and an eRNA (chr11:26,759,157-26,760,455) with the known LM- associated gene *BDNF* [35]; an eRNA (chr12:69,079,117-69,079,445) with the known MM- associated gene *MDM2* [36] (**Fig. 5g**). For each tumor- specific eRNA- regulated target gene set, we calculated GSVA scores for samples within the same tumor type and then stratified them into high, medium, and low score groups. Across different cancer types, we observed significant differential expression of ten hallmark gene sets between the high- and low- score groups (**Fig. 5h**). Further GSEA revealed that cancer-related signaling pathway genes also showed pronounced expression differences between the two groups. Notably, the direction of expression changes for these signaling pathway genes varied by cancer type. For instance, the cell cycle and p53 signaling pathway gene sets were upregulated in the high- score group of AML but downregulated in the high- score group of MM (**Fig. 5j, k**). Similar patterns were recapitulated in mouse samples (**Fig. 5i, Extended Data Fig. 5l, m**). These results underscore the inter-tumor heterogeneity across different cancer types and the complexity of their underlying pathogenesis and disease stages. Moreover, we observed the significant associations between the GSVA scores of tumor-type-specific eRNA- regulated target gene sets and immune-related cell proportions across different cancer samples (**Fig. 5l**). These associations suggest that eRNA-driven transcriptional programs may modulate the composition of the tumor immune microenvironment, potentially influencing immune cell recruitment, activation, or suppression in a cancer-type-specific manner. Collectively, these findings indicate that tumor- specifically activated eRNAs exert tumor- type- dependent functional roles.

### *FOXO1e* is an eRNA cluster regulating t(8;21) AML oncogenesis

To further validate the functional roles of eRNAformer-identified eRNAs in hematological malignancies, we sought to pinpoint eRNA molecules capable of regulating known key oncogenes from a previously constructed eRNA–target network in hematological tumors. Using AML as an example, eRNAformer identified an eRNA cluster, termed *FOXO1e*, which consists of three eRNA loci located approximately 120 kb upstream of *FOXO1*—an oncogene that specifically drives the preleukemic program in t(8;21) AML (**Fig. 6a**). The ENCODE ChIP-seq dataset shows significant enrichment of H3K4me1 and H3K27ac but not H3K4me3 at this region (**Fig. 6a**). Additionally, numerous transcription factor (TF) binding sites and cis-regulatory elements are enriched in this region (**Fig. 6a)**. Moreover, in the t(8;21) AML cell line Kasumi-1, we observed robust enrichment of H3K4me1 and H3K27ac signals, accompanied by the loss of H3K27me3 signal (**Fig. 6b**). In contrast, other cell lines exhibited reduced H3K27ac and increased H3K27me3 signals (**Extended Data Fig. 6a**). These findings indicate that the *FOXO1e* locus adopts an active enhancer state specifically in Kasumi-1 cells. Expression levels of the three eRNAs at the *FOXO1e* locus were highly correlated with *FOXO1* expression in AML RNA-seq samples (**Fig. 6c**). We further examined the expression of these eRNAs across different AML cell lines. Using distinct qPCR primers, we found that all three eRNAs were specifically expressed in Kasumi-1 cells (**Fig. 6d-f**). Similar specific expression was also observed in primary t(8;21) AML patients (**Extended Data Fig. 6b-d**). These results suggest that *FOXO1e* likely exerts a lineage- and context-specific regulatory function in t(8;21) AML cells.

**Fig. 6.**
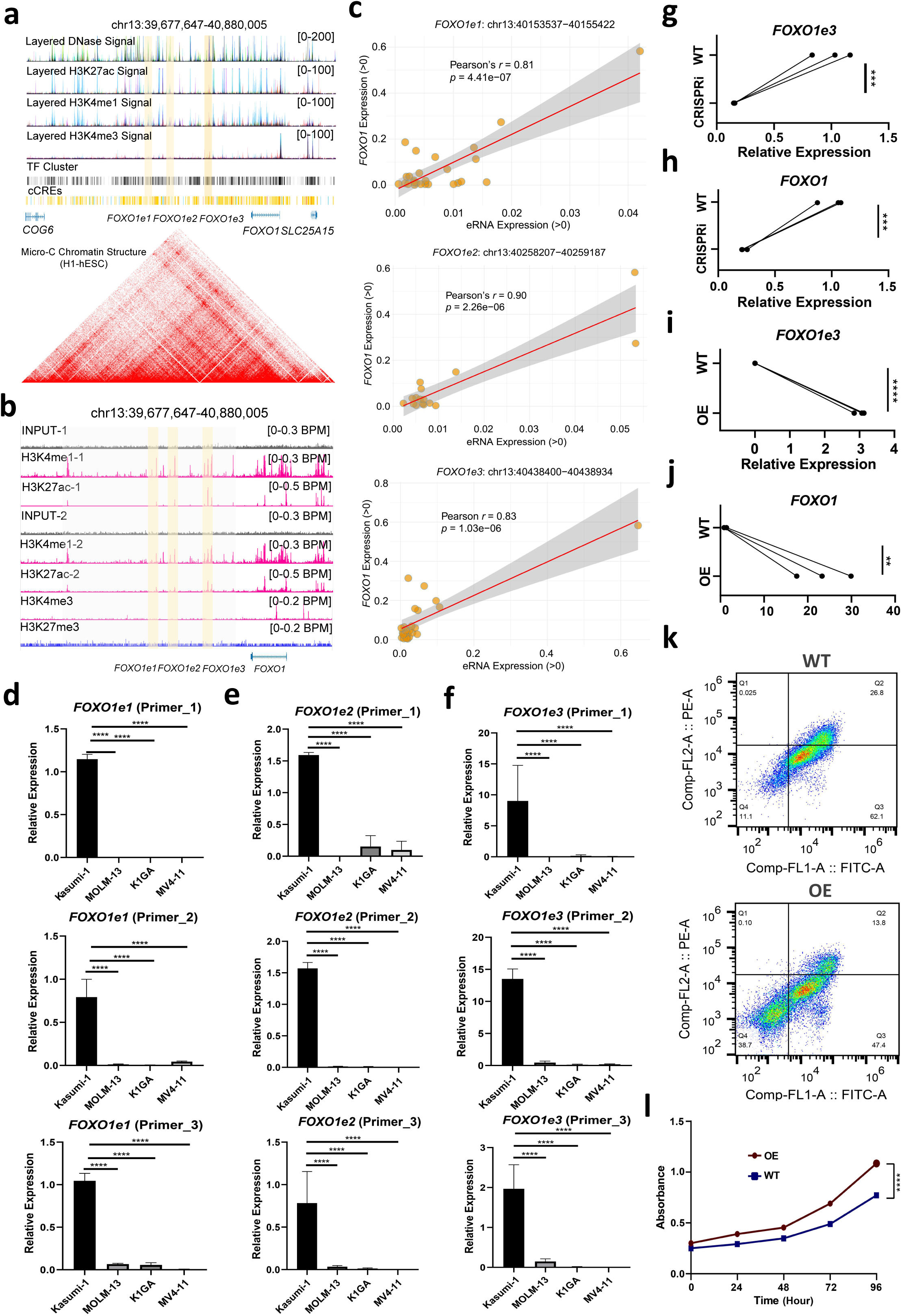
*FOXO1e* is an eRNA cluster that promotes t(8;21) AML oncogenesis. **a)** Genomic context of the *FOXO1e* locus (chr13:39,677,647–40,880,005), including flanking genes, transcriptional orientation, and relative positions of putative regulatory elements. **b)** Enrichment profiles of H3K4me1 (active enhancer mark) and H3K27ac (active promoter/enhancer mark) within and flanking the *FOXO1e* locus, as determined by ChIP-seq. Signal intensities are BPM-based normalization and displayed as read density tracks. **c)** Scatter plot illustrating the correlation between *FOXO1e* and *FOXO1* expression levels, with each point representing an individual sample. Pearson’s correlation coefficient and statistical significance are indicated. The symbol *n* denotes the number of tissue samples used in the correlation analysis. **d)** Quantification of *FOXO1e1* expression levels across a panel of different AML cell lines, assessed by qPCR using three independent primer sets. *p*-value < 0.0001 (****), two-way ANOVA test. **e)** Quantitative PCR analysis of *FOXO1e2* expression in various AML cell lines, performed with three distinct primer pairs. *p*-value < 0.0001 (****), two-way ANOVA test. **f)** Expression profiling of *FOXO1e3* in multiple AML cell lines via qPCR using three different primer sets. *p*-value < 0.0001 (****), two-way ANOVA test. **g)** Dot-line plot demonstrating efficient knockdown of *FOXO1e3* using a CRISPR interference (CRISPRi) system. Each dot represents an independent biological replicate, and lines connect mean values before and after knockdown. *p*-value < 0.0001 (****), two-way ANOVA test. **h)** Dot-line plot showing that *FOXO1e3* knockdown leads to significant downregulation of *FOXO1* mRNA expression. Error bars indicate SEM. *p*-value < 0.05 (*), two-way ANOVA test. **i)** Dot-line plot validating overexpression of *FOXO1e3* using a lentiviral-based expression system. *p*-value < 0.0001 (****), two-way ANOVA test. **j)** Dot-line plot revealing that enforced overexpression of *FOXO1e3* results in marked upregulation of endogenous *FOXO1*. The log-scaled relative expression is shown. *p*-value < 0.0001 (****), two-way ANOVA test. **k)** Flow cytometry analysis demonstrating a marked reduction in apoptosis levels in *FOXO1e3*-overexpressing cells compared to control cells at 96 hours post-transfection. **l)** Dot-line plot showing a significant increase in proliferation levels in *FOXO1e3*-overexpressing cells relative to control cells. *p*-value < 0.0001 (****), two-way ANOVA test.

We next investigated the three-dimensional chromatin architecture surrounding the *FOXO1e* locus. Interestingly, perturbation of *AE* gene expression—a key fusion oncogene driving the preleukemic program in t(8;21) AML—disrupted the spatial chromatin interaction between the *FOXO1e* and *FOXO1* loci, rendering the three-dimensional conformation diffuse and disorganized (**Extended Data Fig. 6e**). This observation supports a model in which *FOXO1e* regulates *FOXO1* expression via chromatin looping. To functionally validate this regulatory relationship, we first used the CRISPRi system to knock down *FOXO1e3*, the most specifically expressed eRNA among the three. *FOXO1e3* knockdown led to a significant reduction in *FOXO1* expression. Furthermore, *FOXO1e3*- knockdown cells exhibited markedly increased apoptosis and decreased proliferation compared to control cells (**Extended Data Fig. 6f-h**). To determine whether *FOXO1e* directly drives *FOXO1* expression as an enhancer RNA, we generated a *FOXO1e3* overexpression cell line. Consistent with the knockdown results, *FOXO1e3* overexpression significantly upregulated *FOXO1* expression, reduced apoptosis, and promoted proliferation in Kasumi-1 cells (**Fig. 6i-l, Extended Data Fig. 6i**). Taken together, these results demonstrate that *FOXO1e* functions as an eRNA cluster that promotes oncogenesis in t(8;21) AML.

## Discussion

Enhancer-derived RNAs (eRNAs) have emerged as critical regulators of gene expression, yet their *de novo* genome-wide detection remains technically challenging. Current methods predominantly rely on sophisticated nascent RNA-seq technologies, which, while precise, are resource-intensive and not readily applicable to large-scale cohorts. In contrast, conventional RNA sequencing (RNA-seq) offers greater cost-effectiveness and experimental accessibility, creating a pressing need for efficient, unbiased computational tools capable of *de novo* mapping of eRNA loci using standard RNA-seq data. To address this gap, we present eRNAformer, a transformer-based multi-modal deep learning framework that integrates DNA sequence and aggregated RNA-seq signals to accurately identify eRNA loci.

Compared with existing methods such as eRNAFinder and DeepITEH, eRNAformer demonstrates superior predictive performance across multiple benchmarking datasets. This advantage stems from two key innovations: its hybrid model architecture and its multimodal data integration strategy. Architecturally, eRNAformer combines convolutional neural networks with transformers in an optimized framework. Parallel multi-scale convolutions capture local sequence features across varying receptive fields, while the transformer encoder models long-range dependencies that are essential for deciphering the regulatory logic of bidirectional transcription. An attention pooling mechanism dynamically weights the most informative regions of input sequences, enabling the model to focus on functionally relevant segments without being constrained by fixed input lengths. Robust regularization techniques—including dropout, batch normalization, and L2 weight decay—enhance generalization. For multimodal integration, eRNAformer incorporates both DNA sequence and RNA-seq signal profiles. This design is grounded in the biological principle that bidirectional eRNA transcription is fundamentally determined by underlying DNA sequences, while aggregated RNA-seq signals provide direct evidence of transcriptional activity. Ablation studies and integrated gradients analysis confirm that the two modalities synergistically enhance prediction performance, with DNA sequence contributions being particularly prominent near transcription start and end sites—regions known to harbor critical regulatory elements for divergent transcription.

By aggregating reads across samples, previous study shows genuine eRNA signals accumulate recurrently, whereas background transcriptional noise tends to occur stochastically and does not consolidate [16]. A distinctive feature of eRNAformer is its use of aggregated RNA-seq reads from multiple samples for eRNA locus assembly. This strategy is particularly important given that eRNAs are typically expressed at low levels in individual samples, leading to limited statistical power for *de novo* detection. Indeed, our analysis demonstrates that bidirectional transcription signals calculated using aggregated samples are markedly improved compared to those derived from individual samples (**Extended Data Fig 1l, m**), validating this design principle.

Beyond methodological advances, our study provides significant biological insights into eRNA function in cancer. Our comprehensive analysis revealed that eRNAformer-identified eRNAs are enriched in evolutionarily constrained regions, associated with complex diseases, and linked to clinically actionable genes and drug responses, highlighting their broad biological and translational relevance. Moreover, applying eRNAformer to hematologic malignancies revealed a comprehensive landscape of cancer-associated eRNAs, leading to the construction of a blood-tumor-specific eRNA database (HMeR). Notably, we identified and experimentally validated *FOXO1e*, a cluster of eRNA loci located approximately 120 kb upstream of *FOXO1*, a known oncogene in acute myeloid leukemia (AML). Through CRISPR activation and inhibition assays, we demonstrated that *FOXO1e* regulates *FOXO1* expression and modulates AML cell proliferation, establishing a functional link between eRNA-mediated regulation and AML pathogenesis. This finding underscores the potential of eRNAs not only as biomarkers but also as therapeutic targets in hematologic malignancies.

Despite its strengths, eRNAformer has limitations that warrant consideration. First, model performance depends on the quality and depth of input RNA-seq data; while aggregation across samples mitigates issues related to low eRNA expression, applications to datasets with limited sample sizes or heterogeneous sample composition may yield reduced sensitivity. Second, although eRNAformer integrates DNA sequence information, it does not explicitly incorporate other regulatory features such as chromatin accessibility, histone modifications, or transcription factor binding profiles, which could further improve predictive accuracy and mechanistic interpretability. Third, the current framework focuses on binary classification of eRNA loci and does not directly model eRNA–target gene interactions or capture the quantitative dynamics of eRNA expression. Additionally, while we functionally validated *FOXO1e* in AML, the vast majority of newly identified eRNAs across other hematologic malignancies remain uncharacterized experimentally, representing a rich resource for future mechanistic studies.

We propose eRNAformer as a valuable computational resource for the research community. By enabling more accurate and comprehensive mapping of eRNA loci, its application to the vast repository of existing and future RNA-seq datasets will facilitate discovery of novel eRNAs and elucidation of their regulatory mechanisms. The blood-cancer-specific eRNA data portal HMeR provide an accessible platform for exploring eRNA expression, regulation, and potential clinical relevance. Looking forward, integrating eRNAformer with single-cell RNA-seq and spatial transcriptomics could further resolve the cell-type-specific and spatial dynamics of eRNA regulation. Extending the framework to incorporate additional modalities such as chromatin conformation data may enable direct prediction of eRNA–target gene interactions, facilitating the identification of functional eRNA–gene circuits in development and disease. Furthermore, the model architecture itself is adaptable and could be applied to other regulatory RNA classification tasks beyond eRNA identification. Collectively, this study establishes eRNAformer as a powerful tool for eRNA discovery and provides a foundation for future investigations into the functional roles of eRNAs in normal physiology and disease.

## Methods

### Curation of benchmark eRNA and non-eRNA loci

To obtain conventional RNA-seq-defined eRNA and non-eRNA loci, we collected 1376 human and 579 mouse ENCODE RNA-seq samples [37] (**Supplementary Dataset 1**). The corresponding Sequence Read Archive (SRA) files were downloaded from the NCBI SRA database. These SRA files were subsequently extracted and converted to FASTQ format using the SRA Toolkit (v3.0.7). For each sample, the downloaded RNA sequencing reads were first trimmed and quality-filtered using Trim Galore (v0.6.5) with the following parameters: -q 20 --phred33 --stringency 3 --trim-n. The high-quality reads were then aligned to the respective reference genome (human GRCh38.p12 or mouse GRCm38.p6 from GENCODE [38]) using HISAT2 (v2.1.0) [39] with the -dta parameter. PCR duplicates were removed using Picard tools (v2.18.2), retaining only uniquely mapped reads. All processed BAM files for human and mouse ENCODE samples were merged into a single species-specific BAM file using SAMtools (v1.15.1) [40], respectively.

Using the merged BAM files, we comprehensively reconstructed transcripts for human and mouse using StringTie (v2.2.1) [41]. StringTie was run with the -m parameter set to 100, ensuring that assembled transcripts were at least 100 bp in length, consistent with the suggested minimum length for eRNAs in a previous study [42]. We implemented a selection procedure to identify transcripts compatible with the known characteristics of eRNAs, including unspliced and bidirectional transcription. First, the transcripts overlapped with known genes (human v28 or mouse vM17 from GENCODE) and multiple-exons transcripts were excluded. Then, single-exon transcripts were selected as candidate eRNA loci. Furthermore, pairs of single-exon transcripts located within 300 base pairs (bp) of each other–a distance suggested by previous studies–were merged as candidate eRNA loci [43–45]. We intersected the genomic coordinates of candidate eRNA loci with known eRNA annotations from FANTOM5 [20] and eRNAbase [21] using BEDTools (v2.31.1). Candidate eRNA loci that completely covered known eRNA annotations were classified as RNA-seq-defined eRNA loci; this was achieved by setting the BEDTools parameter -F to 1.00. In parallel, RNA-seq-defined true non-eRNA loci were derived from assembled multi-exon transcripts. To ensure a balance between positive and negative samples, we selected an equal number of exon loci (>100 bp in length) from these assembled multi-exon transcript loci as RNA-seq-defined non-eRNAs.

### Data preprocess for model

Genomic intervals specifying the chromosomes, start sites, and end sites of benchmarked eRNA and non-eRNA loci were extracted as target regions.

For DNA sequence preprocessing, reference genome sequences (GRCh38.p12 and GRCm38.p6) were retrieved from GENCODE [38]. The FASTA files contain the four standard nucleotides (A, T, C, G) and ‘N’ representing unknown bases. We retained ’N’ nucleotides, encoding all bases using one-hot vectors: A = [1, 0, 0, 0, 0], T = [0, 1, 0, 0, 0], C = [0, 0, 1, 0, 0], G = [0, 0, 0, 1, 0], N = [0, 0, 0, 0, 1].

For RNA-seq input preprocessing, coverage values for eRNA and non-eRNA regions were extracted from ENCODE bigWig files using Python (v.3.9.16) and pyBigWig (v.0.3.22). Coverage values across all ENCODE bigWig files were summed for each nucleotide position. To mitigate extreme value effects, we applied a log(x+1) transformation to the summed coverage.

The model input combines the five-dimensional one-hot encoded DNA sequence and the one-dimensional log-transformed RNA-seq signal into a six-dimensional feature vector per nucleotide position. These vectors were then transposed to form the input tensor for the eRNAformer model.

### Model architecture

eRNAformer takes the DNA sequence and RNA-seq signal as inputs and predicts a binary class label. The architecture consists of an input processing block, a downsampling layer, a positional encoding layer, a transformer tower, an attention pooling layer, and a classification head. This architecture was implemented using PyTorch on servers equipped with NVIDIA Tesla T4 GPUs.

The input processing block is a multi-scale convolutional block that applies four parallel 1D convolutions with different kernel sizes, each with appropriate padding to maintain the sequence length. The outputs of these convolutions are concatenated and passed through batch normalization, a GELU activation, and dropout. This block transforms the input dimensions from (N, C_in, L) to (N, C_out, L), where N is the batch size, C_in is the input dimension, C_out is the hidden dimension, and L is the input sequence length.

The downsampling layer applies a 1D convolution, followed by batch normalization, GELU activation, and dropout. The positional encoding layer adds relative positional information to the sequence. It uses sine and cosine functions of different frequencies to generate positional encodings, which are then added to the input features. A dropout is applied after the addition. The transformer tower is composed of a transformer encoder (using nn.TransformerEncoderLayer) with a hidden dimension and attention heads. Each transformer layer has a feed-forward dimension of 4*hidden_dim and uses GELU activation. Dropout is applied in the attention mechanism and feed-forward network. The attention pooling layer replaces the global average pooling with a learned attention mechanism. It consists of two linear layers with a tanh activation and softmax to produce attention weights for each position. The output is a weighted sum of the transformer outputs across the sequence, resulting in a fixed-size vector per sample. The classification head consists of two linear layers with a RELU activation and dropout in between. The first linear layer reduces the dimension from hidden_dim to hidden_dim/2, and the second produces a single output value. A sigmoid activation is applied to produce the class probability.

### Model hyperparameter optimization

We performed automated hyperparameter optimization for the eRNAformer model using Optuna. For each trial configuration, the model was trained using AdamW optimization with BCE loss while monitoring validation F1 score as the optimization metric. The process explores 13 key hyperparameters (including architecture dimensions and training settings), and implemented epoch-level early stopping when validation performance plateaus. After completing the specified number of trials or timeout duration, the best-performing hyperparameter configuration was saved to a JSON file for subsequent modeling.

### Model training, validation and testing

The data preparation process begins with raw nucleotide sequences and RNA-seq data, which undergo one-hot encoding and log transformation to create combined input tensors of dimension N × 6 × L. The preprocessing pipeline incorporates data augmentation techniques including random reverse-complementation and Gaussian noise injection to RNA-seq channels. The dataset undergoes stratified splitting into training (70%), validation (15%), and test (15%) sets to maintain class balance across partitions. The training process employs the AdamW optimizer. Class weighting automatically balances positive and negative samples based on their dataset representation. The training loop implements automatic mixed precision for computational efficiency and gradient clipping to prevent exploding gradients. A ReduceLROnPlateau scheduler reduces the learning rate by 50% after three epochs without validation improvement. Validation occurs after each training epoch to monitor performance. During validation, the model switches to evaluation mode with gradient computation disabled. The testing phase loads the best-performing model checkpoint based on validation F1 score. Comprehensive evaluation on the training, validation and test sets generates multiple classification metrics, including accuracy, F1 score, AUROC and AUPR. The true label of each locus was utilized for the eRNAformer modeling during the training, validation and test process.

### Performance evaluation and comparison

To systematically evaluate eRNAformer model performance, we employed stratified random splitting experiment, 10-fold cross-validation experiment and multiple external validation experiments. In the random splitting experiment, we conducted 20 random splits of the dataset into training, validation, and test sets and calculated the models’ performance metrics. In the 10-fold cross-validation experiments, we randomly divided the dataset into 10 equal-sized subsets. For each experiment, one subset was used as the validation set, another subset as the test set, and the remaining subsets as the training set. We conducted 10 experiments to ensure that each subset served as the validation set and as the test set exactly once. For external validations, we recursively evaluated the predictive performance of each trained model on another benchmark dataset. To benchmark the predictive capability of eRNAformer, we performed a comprehensive comparison of eRNAformer with eRNAFinder [18] and DeepITEH [17] using the same evaluation experiments above. The FANTOM5-derived benchmarking dataset was employed for training the model to mitigate concerns regarding the potential presence of false-positive eRNA locus annotations in the eRNAbase database.

eRNAFinder takes the transcription signal profiles as input, whereas DeepITEH takes sequence and epigenetic features as input. We strictly followed the same data pre-processing as the original studies for these two methods. For eRNAFinder, signal profiles of transcription were calculated using deepTools (v3.5.3). Each genomic region (eRNA or non-eRNA locus) was standardized to a fixed length of 900 bp. This region was then divided into 90 equally sized bins (10 bp each). The Bins per Million mapped reads (BPM) signal intensity was computed for each bin. The aggregated signal per bin was defined as the mean signal intensity across all relevant ENCODE samples. For each region, the resulting 90 aggregated signal values per transcription signal served for *P_bt_* calculation. eRNAFinder did not need to be trained and the predictive performance were calculated using the list of *P_bt_* as described in our previous work [18]. For DeepITEH, we did not determine the location of sequence and epigenomic features as the eRNA loci are explicitly defined in this study. We computed epigenomic features using data from ENCODE for five histone marks (H3K4me1, H3K27ac, H3K9ac, H3K4me3, and H3K27me3). The histone modification level for each genomic interval was defined by its mean signal intensity across all samples, which we obtained using UCSC bigWigAverageOverBed tool. According to DeepITEH architecture, we used a bidirectional LSTM to encode DNA sequences and integrated them with histone features through a fusion network. The hyperparameters and performances of DeepITEH models were optimized and calculated using the same method as eRNAformer.

### Feature contribution calculation

To interpret the predictions of our eRNAformer model and deconstruct the relative importance of its multimodal inputs, we employed a feature attribution analysis based on the Integrated Gradients method. Our approach quantitatively attributes the model’s output for a given genomic interval to each individual input feature by integrating the model’s gradients along a straight-line path from a defined baseline input to the actual input sequence. We defined the baseline using a neutral reference state, where the DNA sequence channels were set to a uniform distribution (0.2 probability for each nucleotide) and the RNA signal channel was set to zero, representing the log-transformed value of a zero signal. For each interval, the computed attribution scores were systematically aggregated across the five DNA one-hot-encoded channels and the single RNA signal channel to produce two distinct contribution scores: DNA contribution and RNA contribution. The analysis was performed on a held-out set of genomic intervals. The resulting attribution maps for each interval were saved, and a comprehensive global analysis was conducted by summing absolute contribution scores across all evaluated intervals. This allowed us to determine the total contribution of each modality—DNA and RNA—to the model’s predictive output. Furthermore, we analyzed the positional dependence of these contributions by normalizing each interval to a fixed number of bins (1000) and averaging attribution scores at each relative position across all intervals, providing a genome-wide view of the salient features learned by the model.

### ENCODE H3K4me1 and H3K27ac ChIP-seq data analysis

ChIP-seq signals of H3K4me1 and H3K27ac at eRNAformer-identified human and mouse eRNA loci were extracted from ENCODE BigWig files using the UCSC tool bigWigAverageOverBed (v1.04.00). We included all files that were in BigWig format, aligned to the hg38 or mm10 genome assemblies, and represented fold-change over control signals. The “mean” column from the output was used to quantify signal intensity. Signals were profiled across the entire length of enhancers. Additionally, for eRNA loci of interest, we sampled signals in 50-bp non-overlapping windows across a 1 kb flanking region (1 kb upstream and downstream). The resulting 41 data points (comprising the central element and 20 flanking windows on each side) were converted to Z-scores. Meta-profiles of chromatin features were then generated by averaging the Z-scores across all aligned sequences.

### Evolutionary constraints and disease association analysis

For each eRNA locus, we extracted flanking regions of equal length upstream and downstream. We then downloaded chromosome-specific VCF files from [gnomAD](https://gnomad.broadinstitute.org/data) and extracted genetic variants located within the eRNA loci and their flanking regions. The variants were grouped based on their Minor Allele Frequency (MAF) to compare the distribution of variant counts across the different regions. Furthermore, to investigate the evolutionary conservation of these variants across species, we obtained the bigwig file of PhyloP conservation score profile across human hg38 reference genome from [UCSC Genome Browser](https://hgdownload.soe.ucsc.edu/goldenPath/hg38/phastCons100way/). The PhyloP conservation score of each variant was extracted using bigWigAverageOverBed tool.

To investigate the associations between enhancer RNAs (eRNAs) and human diseases, we first identified putative target genes located within 1 megabase (Mb) of each eRNA using the GENCODE v28 annotation file in GFF3 format. Disease-related information for these eRNAs and their target genes was then extracted by cross-referencing with clinically relevant gene variants from the ClinVar database, using its provided variant call format (VCF) files. For expression analysis, we retrieved RNA-seq data from the GTEx project via the recount3 R package [46], downloading bigWig files representing expression signals. Expression levels of eRNA loci and their corresponding target genes across different GTEx samples were quantified using the pyBigWig package (v0.3.0). To prevent ambiguity in expression quantification, our analysis was confined to eRNA loci located outside of known boundaries of spliced target genes, including exons, UTRs, and transcription start/end sites. In cases where an eRNA locus partially overlaps a known spliced gene, the expression value derived from its non-overlapping region was used to represent the locus’s expression.

We employed two methods to calculate expression correlation: within-tissue correlation was computed based on the expression values of eRNAs and target genes across samples from the same tissue, while inter-tissue correlation was calculated using the average expression values of eRNAs and target genes from merged samples of the same tissue type. For exploring potential regulatory mechanisms, we downloaded publicly available eQTL summary statistics from the GTEx portal (https://gtexportal.org/). By mapping these eQTLs to genomic coordinates, we examined whether any eQTLs associated with the target genes overlapped with the eRNA loci, thereby exploring potential regulatory mechanisms.

### Pan-cancer analysis of eRNAformer-identified eRNAs

The RNA-seq bigWig files of 33 types of TCGA cancers were downloaded using recount3 R package [46]. Cancer signaling pathway genes were extracted from KEGG database [47]. Clinically actionable genes and immune checkpoints were obtained from a previous publication [7]. Gene expression and drug sensitivity datasets of cancer cell lines were downloaded from the CCLE [48], CTRP [49] and GDSC [50]. We applied the same method used for GTEx samples to quantify the expression of eRNA loci and their target genes in the TCGA cohort. The TF binding motifs were collected from JASPAR database [51]. TF binding motif searching of eRNA loci was performed for each cluster using MEME Suite (v5.4.1) Analysis of Motif Enrichment (AME)” program [52]. We identified putative eRNA regulators as transcription factors (TFs) significantly correlated with an individual eRNA (Pearson’s *r* ≥ 0.7, false discovery rate [FDR] < 0.01) or showing both correlation (*r* > 0) and a binding motif in a given cancer. We identified eRNA putative target genes based on close distance (≤ 1MB) and co-expression (r ≥ 0.3 and FDR < 0.05) between individual eRNAs and their putative target genes in each cancer type. The human Hi-C datasets from ENCODE were used for 3-dimensional chromatin interaction analysis. To analyze the 3D chromatin interaction, the downloaded .hic files were converted and merged to Hi-C contact matrices at 10kb resolution using a custom script. The binned raw Hi-C contact matrices were corrected by Knight–Ruiz matrix balancing method using “hicCorrectMatrix” command in HiCExplorer tool [53]. The corrected Hi-C contact matrices were further converted to Hi-C Observed/Expected (O/E) matrix using HiCExplorer “hicTransform” command. Randomly selected and distance-matched intra-chromosome genomic region pairs were produced as control when comparing with eRNA-target gene links. The custom R codes were used to analyze and visualize the Hi-C interaction for eRNAs-target gene links and randomly selected pairs. RNA-seq data of Vorinostat treatment were downloaded from GSE110948. Using the same preprocess procedure as described above, the RNA-seq samples were quality filtered and aligned human hg38 reference genome. The, we converted the aligned BAM file to to bigWig file and calculated the expression signal value of eRNAformer-identified eRNA loci using bamCoverage command from deepTools and pyBigWig (v.0.3.22). We selected the eRNAs with absolute log2(fold change) larger than 0.5 as differentially expressed eRNAs.

### *De novo* mapping eRNAs in hematological malignancies

To systematically identify undiscovered hematological malignancies-specific eRNAs, we downloaded collected 1497 conventional RNA-seq samples of human hematological malignancies (AML: 490, CML: 650, LM: 157 and MM: 200) and 1168 conventional RNA-seq samples of mouse hematological malignancies (AML: 297, CML: 570, LM: 101 and MM: 200) (**Supplementary Dataset 5**). Following the RNA-seq data processing steps previously described, we aligned the samples to the reference genome. Alignment results from samples of the same tumor type were then merged into single BAM files for transcript assembly. From the assembled transcripts, we selected unannotated single-exon transcripts that did not overlap any annotated genes. Transcripts located within 300 bp of each other were merged to define candidate eRNA loci. To accurately predict hematological malignancies-specific eRNAs, we used the model pre-trained on ENCODE RNA-seq data as the foundation. We constructed a new training dataset comprising: 1) candidate eRNA loci in hematological malignancies that fully overlapped known eRNA annotations from either FANTOM5 or eRNAbase (serving as positive eRNA loci), and 2) an equal number of exonic regions randomly selected from assembled multi-exon transcripts (serving as RNA-seq-defined non-eRNAs). This dataset was used to fine-tune the pre-trained model. The fine-tuned model was then applied to predict eRNAs among the remaining unannotated candidate loci.

### Hematological malignancies eRNA analysis

We applied the same method used for TCGA samples to quantify the expression of eRNA loci in the hematological malignancy samples. The TF- eRNA–target network for each hematological malignancy was constructed using the same approach as in the pan- cancer analysis. We identified eRNAs and their target genes that are consistently activated across multiple hematological malignancies, defined as those with a mean expression level higher in tumor samples than in normal samples and a standard deviation of expression changes among tumor types less than 0.5. Cancer-type specific eRNAs and target genes were identified using the Jensen–Shannon (JS) divergence score [54]; eRNAs with a JS score greater than 0.4 were defined as specifically expressed in a given cancer type. The GSVA scores of eRNA target gene sets were calculated using the GSVA R package (v1.52.3) with default parameters. Gene set enrichment analysis (GSEA) comparing low and high GSVA groups was performed using the fgsea R package (v1.30.0) with default parameters.

### HMeR Data portal

The data portal was built based on Spring Boot (v2.7.18) (https://spring.io/projects/spring-boot) and MySQL (v8.0.33) (https://www.mysql.com).The frontend interface was created by HTML5, Tailwind CSS (v3.4.1) (https://tailwindcss.com), ECharts (v5.4.3) (https://echarts.apache.org), and native JavaScript (ES6). Data query and rendering were implemented using JdbcTemplate in the backend.

### AML ChIP-seq data analysis

The ChIP sequencing reads of H3K4me1 and H3K27ac were downloaded, extracted, trimmed and quality-filtered using the same method as RNA-seq data pre-processing. We mapped the high-quality sequencing reads to mouse mm10 reference genome using Bowtie2 (v2.5.1). For single-end sequencing reads, the Bowtie2 parameters for H3K4me1 and H3K27ac ChIP-seq data were “-t -q -N 1 -L 25 -X 2000 --no-mixed --no-discordant”. For paired-end sequencing reads, the Bowtie2 parameters for H3K4me1 and H3K27ac ChIP-seq data were “-t -q -N 1 -L 25 --no-mixed --no-discordant”. PCR duplicates were removed. The BAM files of the same developmental stage were merged, sorted, and indexed for further analysis With SAMtools (v1.15.1). The “bamCoverage” and “bamCompare” commands from deepTools (v3.5.3) were employed for downstream analysis. Using “bamCoverage” command with the parameter “-normalizeUsing BPM -of bigwig -binSize 10, the raw reads signal was normalized to Bins per Million mapped reads (BPM) signal and converted to bigwig signal files. For ChIP-seq data with input sequencing control, the ChIP-seq signal was corrected using log2 ratio transformation between IP signal and input signal by “bamCompare” command. The “computeMatrix”, “plotProfile” and “plotHeatmap” commands of deepTools were used to produce the ChIP signal curve for the targeted DNA regions.

### AML capture Hi-C data analysis

Capture Hi- C data were processed using HiC-Pro (v3.1.0) [55]. Paired- end reads were aligned to the human reference genome with Bowtie2. Reads were assigned to restriction fragments. Valid interaction pairs (non- self, non-PCR duplicate) were retained. Capture targets were provided as a BED file to focus the analysis on regions of interest. Raw contact matrices were built at 10 kb resolutions. The binned raw Hi-C contact matrices were Knight–Ruiz corrected and further converted to Hi-C Observed/Expected (O/E) matrix using HiCExplorer (v3.7.2) [53]. The corrected Hi-C contact matrices were further converted to Hi-C Observed/Expected (O/E) matrix using HiCExplorer “hicTransform” command. The custom R codes were used to visualize the Hi-C interaction.

### AML cell lines

The Kasumi-1 cell line was cultured in RPMI 1640 medium supplemented with 20% fetal bovine serum (FBS) and 1% penicillin/streptomycin (P/S). The 293T cell line was cultured in DMEM supplemented with 10% fetal bovine serum (FBS) and 1% penicillin/streptomycin (P/S). To maintain optimal growth conditions and prevent contamination, the culture medium was replaced twice weekly. When cell confluence reached approximately 80%, cells were passaged at a 1:3 ratio to ensure continuous proliferation and facilitate subsequent experiments.

### CRISPR/dCas9-KRAB

Lentiviral particles were produced using the following plasmids: Lenti dCas9-KRAB and lenti-sgRNA (MS2) puro-optimized backbone (#73797). HEK293T cells (passages 2–5) were seeded in 10 cm dishes and transfected at 80–90% confluency. Transfection was performed using a mass ratio of PSPA: PMD2G: target plasmid = 3:2:5 (total 15 μg DNA per dish). Polyethylenimine (PEI) and DNA were mixed at a volume-to-mass ratio of 4:1 (PEI: DNA) in Opti-MEM to a final volume of 1 mL. The mixture was incubated for 15 min at room temperature before being added to the cells. Viral supernatants were collected at 48 h and 72 h post-transfection, centrifuged at 4000 rpm for 15 min to remove debris, filtered through a 0.45 μm PES membrane, aliquoted, and stored at −80 °C (avoiding repeated freeze-thaw cycles). Kasumi-1 cells in the logarithmic growth phase were seeded and transduced using Lenti dCas9-KRAB lentivirus. Transduced cells were selected with blasticidin (30 μg/mL) for 10–14 days to obtain the dCas9-KRAB stable pool. This pool was then transduced with sgRNA lentivirus and selected with puromycin (5 μg/mL) to establish the final CRISPRi stable cell line. gRNA sequences were provided in **Supplementary Dataset S6**.

### Overexpression

Overexpression plasmids were constructed, transformed into competent cells, cultured, and extracted after monoclonal picking. Lentivirus was packaged in 293T cells (70%-80% confluence): DMEM-diluted PEI and plasmids were mixed, incubated, added to cells, and viral supernatant was collected at 48/72 h, centrifuged, filtered and stored. Kasumi-1 cells were transduced with lentivirus (MOI 10/100) plus Polybrene, centrifuged, re-infected, cultured, and screened with puromycin to obtain stable cell lines.

### RT-qPCR

Total RNA was extracted from cells using Servicebio RNA extraction kit after PBS washing, eluted with RNase-free water, and qualified samples were stored at −80 °C. Reverse transcription was performed with Takara Two-Step RT Kit: genomic DNA was eliminated first, then 5X RT Premix was added for reaction at 37 °C for 10 min and 85 °C for 5 sec. qPCR was conducted with SYBR Green qPCR Mix, three technical replicates per sample; amplification included pre-denaturation (95 °C, 3min), 40 cycles (95 °C 5s, 60 °C 34s) and melting curve analysis. Relative mRNA expression was calculated by 2^(-ΔΔCt) method. Primer sequences were provided in **Supplementary Dataset S6**.

### Cell proliferation assay

Cell proliferation was assessed using the Cell Counting Kit-8 (CCK-8) assay. Briefly, cells in the logarithmic growth phase were collected. 5×10⁹ cells were added to each well. At 0 h, 24 h, 48 h and 96 h, 10 μL of CCK-8 solution was added to each well, and the plate was incubated in the incubator away from light for 3-4 h. The optical density (OD) value at 450 nm was measured using a microplate reader.

### Cell apoptosis assay

Cells in logarithmic growth phase were collected, centrifuged (600 rpm, 5 min), resuspended, adjusted to 3×10⁵ cells/mL, seeded into 24-well plates, and harvested at 24, 48, 72 h, 96h. Harvested cells were centrifuged (1500 rpm, 5 min, 4 °C), washed with pre-cooled 1×PBS, resuspended in 200 μL 1×Binding Buffer, stained with 2 μL Annexin V-APC and 1 μL PI away from light for 5 min, then detected by flow cytometry. Apoptosis rate was compared by Annexin V-positive cell proportion.

### Quantification, statistical analysis and visualization

All NGS data quantification and statistical analyses were performed using R (v4.4.1). All NGS results visualization were performed using R (v4.4.1) and Integrative Genomics Viewer (v2.19.1). Other data quantification, statistical analyses and visualization were conducted with GraphPad Prism 8 software. Details of specific statistical tests are detailed in the figure legends.

## Code availability

The code of eRNAformer has been deposited at Github and is publicly available at https://github.com/tkrwy/eRNAformer.

## Data availability

All data are available within the article and its **Supplementary Datasets**. Accession codes of the published data in GEO used in this study are as follows: AML H3K4me1 and H3K27ac, GSE115115 and GSE128528; AML H3K4me3 and H3K27me3, [GSE62847](https://www.ncbi.nlm.nih.gov/geo/query/acc.cgi?acc=GSE128528) and GSE128528; AML capture Hi-C, GSE117107. The expression of blood-cancer-specific eRNAs, TF-eRNA and eRNA-target links are available to query and visualize at our online data portal.

