## Extended Data for "eRNAformer enables genome-wide *de novo* mapping of enhancer-derived RNA loci"

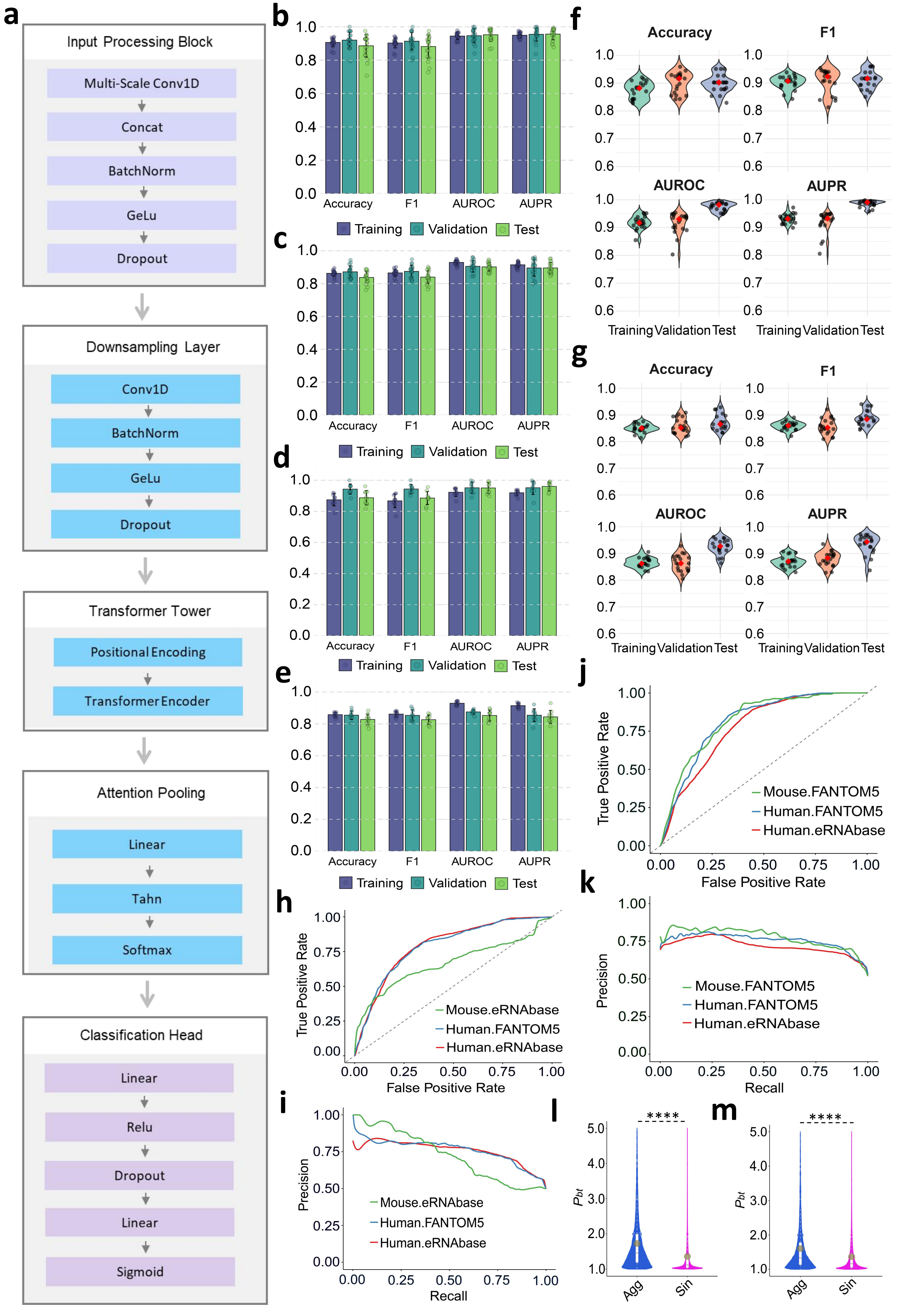
a
Input Processing Block
Multi-Scale Conv1D
Concat
BatchNorm
GeLu
Dropout
Downsampling Layer
Conv1D
BatchNorm
GeLu
Dropout
Transformer Tower
Positional Encoding
Transformer Encoder
Attention Pooling
Linear
Tahn
Softmax
Classification Head
Linear
Relu
Dropout
Linear
Sigmoid
b
c
d
e
Accuracy
F1
AUROC
AUPR
Training
Validation
Test
f
g
Accuracy
F1
AUROC
AUPR
Training Validation Test
h
True Positive Rate
False Positive Rate
Mouse.eRNAbase
Human.FANTOM5
Human.eRNAbase
i
Precision
Recall
Mouse.eRNAbase
Human.FANTOM5
Human.eRNAbase
j
True Positive Rate
False Positive Rate
Mouse.FANTOM5
Human.FANTOM5
Human.eRNAbase
k
Precision
Recall
Mouse.FANTOM5
Human.FANTOM5
Human.eRNAbase
l
m
****
$P_{bt}$
Agg
Sin

**Extended Data Fig. S1**. Model design and performance of eRNAformer, related to **Fig. 1**.
**a)** Detailed architecture of different eRNAformer layers.
**b)** Prediction performance of the eRNAformer model constructed on the mouse FANTOM5-derived benchmark dataset across 20 random data splits.
**c)** Prediction performance of the eRNAformer model constructed on the mouse eRNAbase-derived benchmark dataset across 20 random data splits.
**d)** Prediction performance of the eRNAformer model constructed on the mouse FANTOM5-derived benchmark dataset across10-fold cross-validation experiments.
**e)** Prediction performance of the eRNAformer model constructed on the mouse eRNAbase-derived benchmark dataset across 10-fold cross-validation experiments.
**f)** Prediction performance of the eRNAformer model constructed on the mouse FANTOM5-derived benchmark dataset across 20 independently generated non-eRNA loci sets.
**g)** Prediction performance of the eRNAformer model constructed on the mouse eRNAbase-derived benchmark dataset across 20 independently generated non-eRNA loci sets.
**h)** ROC curves of the eRNAformer model constructed on the mouse FANTOM5-derived benchmark dataset for intra- and inter-species external data predictions.
**i)** PR curves of the eRNAformer model constructed on the mouse FANTOM5-derived benchmark dataset for intra- and inter-species external data predictions.
**j)** ROC curves of the eRNAformer model constructed on the mouse eRNAbase-derived benchmark dataset for intra- and inter-species external data predictions.
**k)** PR curves of the eRNAformer model constructed on the mouse eRNAbase-derived benchmark dataset for intra- and inter-species external data predictions.
**l)** Violin plots showing the *Pbt* distribution of known eRNA loci in human genome calculated by aggregated and single RNA-seq samples. *p*-value < 0.0001 (****), two-sided Wilcoxon signed-rank test. Agg, Aggregated RNA-seq samples; Sin, Single RNA-seq samples.
**m)** Violin plots showing the *Pbt* distribution of known eRNA loci in mouse genome calculated by aggregated and single RNA-seq samples. *p*-value < 0.0001 (****), two-sided Wilcoxon signed-rank test. Agg, Aggregated RNA-seq samples; Sin, Single RNA-seq samples.

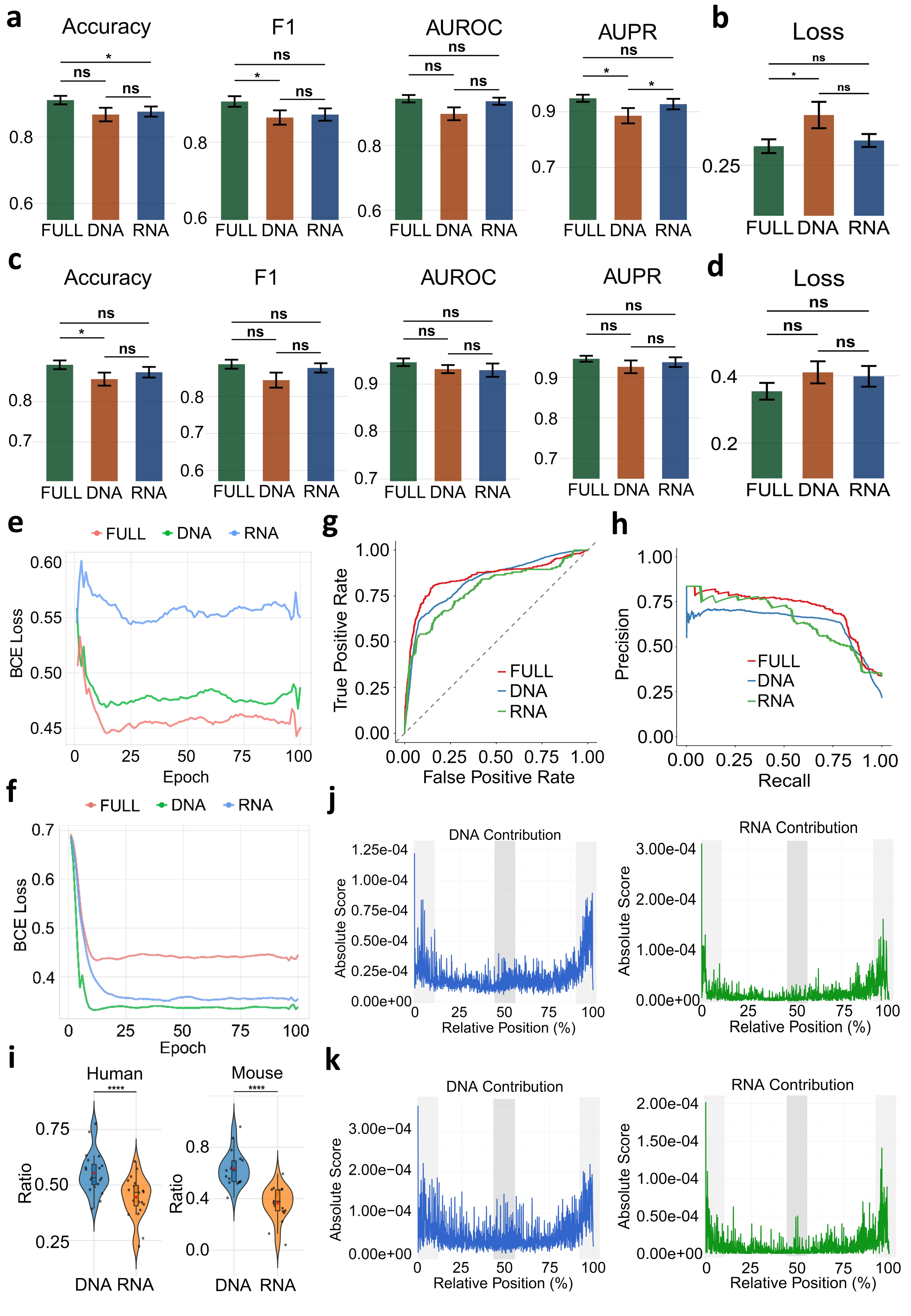

a
Accuracy
F1
AUROC
AUPR
b
Loss
ns
*
FULL DNA RNA
c
d
e
FULL DNA RNA
BCE Loss
Epoch
f
g
True Positive Rate
False Positive Rate
h
Precision
Recall
i
Human
Mouse
****
Ratio
j
DNA Contribution
RNA Contribution
Absolute Score
Relative Position (%)
k

**Extended Data Fig. S2**. Combining DNA sequence and RNA-seq data synergistically improves the prediction of eRNA loci, related to **Fig. 2**.

**a)** Performance comparison between the full model and single-input models trained from mouse FANTOM5-derived benchmark dataset on the validation set across 20 random data splits; $p$-value < 0.05 (*), $p$-value ≥ 0.05 (ns), two-sided Wilcoxon signed-rank test.

**b)** BCE loss comparison between the full model and single-input models trained from mouse FANTOM5-derived benchmark dataset on the validation set across 20 random data splits; $p$-value < 0.05 (*), $p$-value ≥ 0.05 (ns), two-sided Wilcoxon signed-rank test.

**c)** Performance comparison between the full model and single-input models trained from mouse FANTOM5-derived benchmark dataset on the test set across 20 random data splits; $p$-value < 0.05 (*), $p$-value ≥ 0.05 (ns), two-sided Wilcoxon signed-rank test.

**d)** BCE loss comparison between the full model and single-input models trained from mouse FANTOM5-derived benchmark dataset on the mouse test set across 20 random data splits; $p$-value ≥ 0.05 (ns), two-sided Wilcoxon signed-rank test.

**e)** BCE loss curves of the full model and single-input models at 100 epoches in human validation dataset.

**f)** BCE loss curves of the full model and single-input models at 100 epoches in mouse validation dataset.

**g)** ROC curves of the full model and single-input models, trained on the mouse FANTOM5-derived benchmark dataset, when predicting on the human FANTOM5-derived benchmark dataset.

**h)** PR curves of the full model and single-input models, trained on the mouse FANTOM5-derived benchmark dataset, when predicting on the human FANTOM5-derived benchmark dataset.

**i)** DNA and RNA contribution of human and mouse models across 20 random data splits; $p$-value < 0.0001 (****), two-sided Wilcoxon signed-rank test.

**j)** Mean DNA and RNA contribution profile across all eRNA and non-eRNA loci for full model trained from human FANTOM5-derived benchmark dataset.

**k)** Mean DNA and RNA contribution profile across all eRNA and non-eRNA loci for full model trained from mouse FANTOM5-derived benchmark dataset.

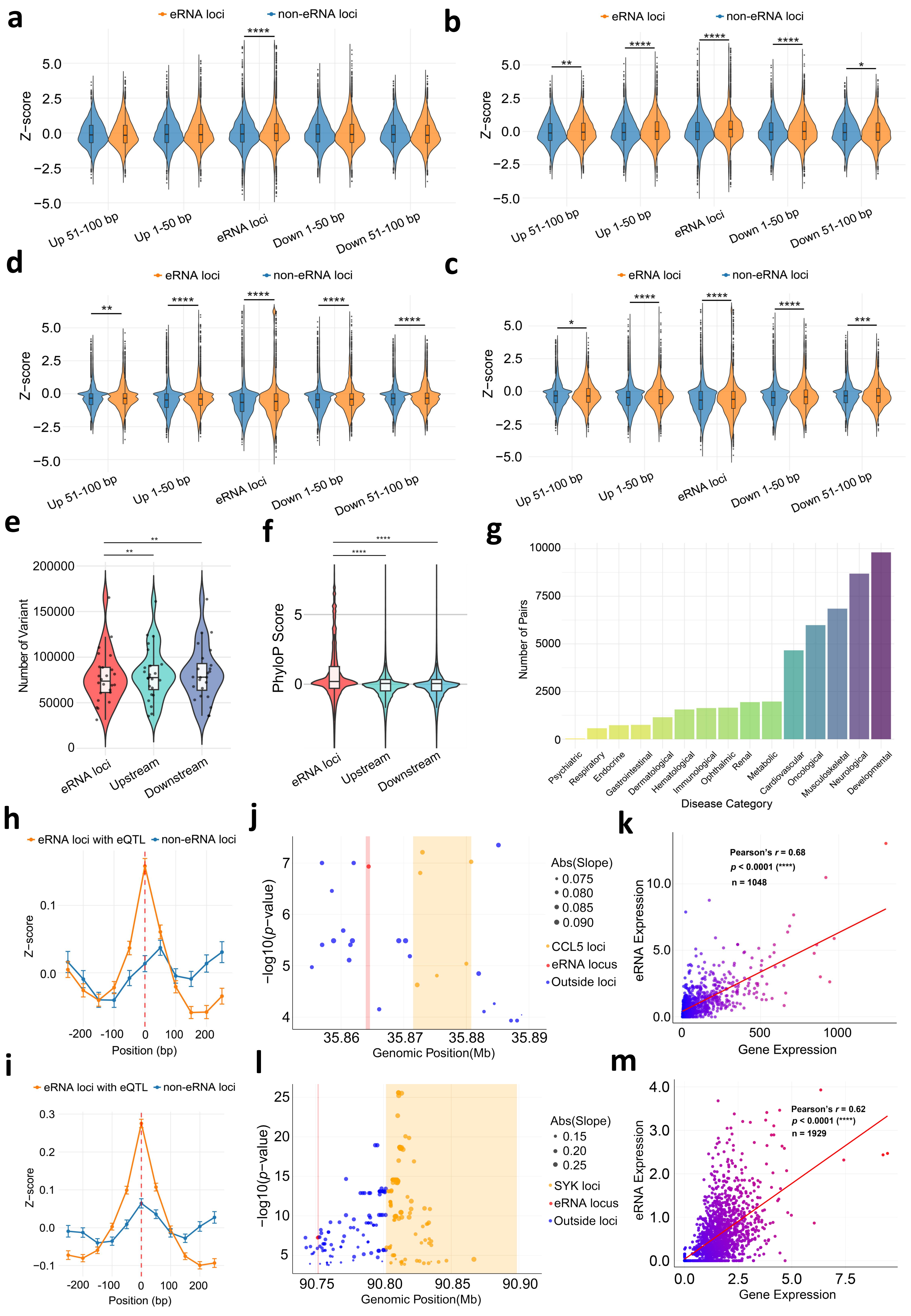

a
eRNA loci
non-eRNA loci
****
Z−score
5.0
2.5
0.0
−2.5
−5.0
Up 51-100 bp
Up 1-50 bp
eRNA loci
Down 1-50 bp
Down 51-100 bp
b
eRNA loci
non-eRNA loci
**
****
****
****
*
Z−score
Up 51-100 bp
Up 1-50 bp
eRNA loci
Down 1-50 bp
Down 51-100 bp
d
eRNA loci
non-eRNA loci
**
****
****
****
****
Z−score
Up 51-100 bp
Up 1-50 bp
eRNA loci
Down 1-50 bp
Down 51-100 bp
c
eRNA loci
non-eRNA loci
*
****
****
****
***
Z−score
Up 51-100 bp
Up 1-50 bp
eRNA loci
Down 1-50 bp
Down 51-100 bp
e
**
**
Number of Variant
200000
150000
100000
50000
0
eRNA loci
Upstream
Downstream
f
****
****
PhyloP Score
5
0
eRNA loci
Upstream
Downstream
g
Number of Pairs
10000
7500
5000
2500
0
Psychiatric
Respiratory
Endocrine
Gastrointestinal
Dermatological
Hematological
Immunological
Ophthalmic
Renal
Metabolic
Cardiovascular
Oncological
Musculoskeletal
Neurological
Developmental
Disease Category
h
eRNA loci with eQTL
non-eRNA loci
Z−score
0.1
0.0
-200
-100
0
100
200
Position (bp)
j
−log10(p−value)
7
6
5
4
35.86
35.87
35.88
35.89
Genomic Position(Mb)
Abs(Slope)
0.075
0.080
0.085
0.090
CCL5 loci
eRNA locus
Outside loci
k
Pearson's r = 0.68
p < 0.0001 (****)
n = 1048
eRNA Expression
10.0
5.0
0.0
0
500
1000
Gene Expression
i
eRNA loci with eQTL
non-eRNA loci
Z−score
0.3
0.2
0.1
0.0
−0.1
-200
-100
0
100
200
Position (bp)
l
−log10(p−value)
25
20
15
10
5
90.75
90.80
90.85
90.90
Genomic Position(Mb)
Abs(Slope)
0.15
0.20
0.25
SYK loci
eRNA locus
Outside loci
m
Pearson's r = 0.62
p < 0.0001 (****)
n = 1929
eRNA Expression
4.0
3.0
2.0
1.0
0.0
0.0
2.5
5.0
7.5
Gene Expression

**Extended Data Fig. S3**. eRNAformer maps evolutionarily constrained eRNAs associated with complex diseases, related to **Fig. 3**.
**a)** Violin plots with boxplots showing the Z-score normalized H3K4me1 signals at human eRNA loci identified by eRNAformer using aggregated ENCODE RNA-seq data, compared to assembled single-exon non-eRNA loci and their flanking 200 bp regions; *p*-value < 0.0001 (****), two-sided Mann–Whitney U-test.
**b)** Violin plots with boxplots displaying the Z-score normalized H3K27ac signals at human eRNA loci identified by eRNAformer using aggregated ENCODE RNA-seq data, compared to assembled single-exon non-eRNA loci and their flanking 200 bp regions; *p*-value < 0.0001 (****), *p*-value < 0.01 (**), *p*-value < 0.05 (*), two-sided Mann–Whitney U-test.
**c)** Violin plots with boxplots illustrating the Z-score normalized H3K4me1 signals at mouse eRNA loci identified by eRNAformer using aggregated ENCODE RNA-seq data, compared to assembled single-exon non-eRNA loci and their flanking 200 bp regions; *p*-value < 0.0001 (****), *p*-value < 0.01 (**), two-sided Mann–Whitney U-test.
**d)** Violin plots with boxplots depicting the Z-score normalized H3K27ac signals at mouse eRNA loci identified by eRNAformer using aggregated ENCODE RNA-seq data, compared to assembled single-exon non-eRNA loci and their flanking 200 bp regions; *p*-value < 0.0001 (****), *p*-value < 0.001 (***), *p*-value < 0.05 (*), two-sided Mann–Whitney U-test.
**e)** Violin plots with boxplots presenting the distribution of variant counts in eRNA loci predicted by eRNAformer and in flanking regions of equal length; *p*-value < 0.01 (**), two-sided Wilcoxon signed-rank test. Each point corresponds to the number of variants on a single chromosome.
**f)** Violin plots with boxplots showing the distribution of PhyloP conservation scores for variants located within eRNAformer-predicted eRNA loci and comparable flanking regions; *p*-value < 0.0001 (****), two-sided Mann–Whitney U-test.
**g)** Bar plot showing the number of eRNA-target gene pairs associated with different disease types that exhibit significant correlations, defined as a Pearson correlation coefficient > 0.6 and a p-value < 0.01.
**h)** Line plots with error bars showing Z-score-normalized H3K4me1 signals at human eRNA loci harboring eQTLs identified by eRNAformer using aggregated ENCODE RNA-seq data, compared to those at assembled non-eRNA loci and their flanking 200 bp regions.
**i)** Line plots with error bars showing Z-score-normalized H3K27ac signals at human eRNA loci harboring eQTLs identified by eRNAformer using aggregated ENCODE RNA-seq data, compared to those at assembled non-eRNA loci and their flanking 200 bp regions.
**j)** Scatter plot illustrating an eQTL of the target gene CCL5 located within the eRNA region (chr17:35,863,936-35,864,615). The area shaded in red denotes the eRNA locus, while the area in yellow corresponds to the gene region.
**k)** Scatter plot displaying the expression correlation between the eRNA (chr17:35,863,936-35,864,615) and its target gene CCL5 in whole blood tissue. The symbol *n* denotes the number of whole-blood samples used in the correlation analysis.
**l)** Scatter plot showing an eQTL of the target gene SYK located within the eRNA region (chr9:90,751,240-90,752,230). The area shaded in red denotes the eRNA locus, while the area in yellow corresponds to the gene region.
**m)** Scatter plot depicting the expression correlation between the eRNA (chr9:90,751,240-90,752,230) and its target gene SYK in skin tissue. The symbol *n* denotes the number of whole-blood samples used in the correlation analysis.

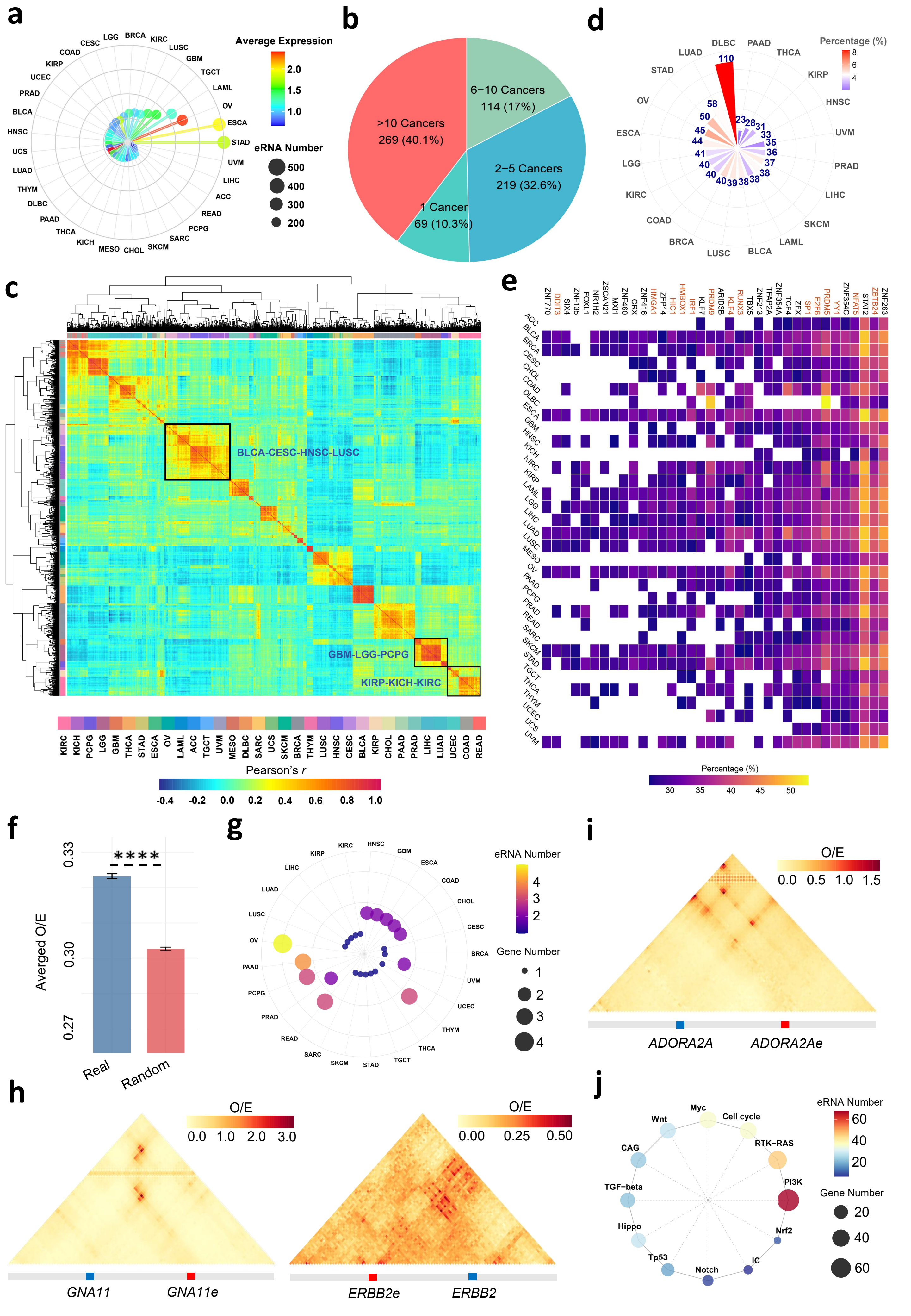
a
Average Expression
2.0
1.5
1.0
eRNA Number
500
400
300
200
LGG BRCA KIRC LUSC GBM TGCT LAML OV ESCA STAD UVM LIHC ACC READ PCPG SARC SKCM CHOL MESO KICH THCA PAAD DLBC THYM LUAD UCS HNSC BLCA PRAD UCEC KIRP COAD CESC
b
>10 Cancers
269 (40.1%)
6−10 Cancers
114 (17%)
2−5 Cancers
219 (32.6%)
1 Cancer
69 (10.3%)
d
Percentage (%)
8
6
4
DLBC PAAD THCA KIRP HNSC UVM PRAD LIHC SKCM LAML BLCA LUSC BRCA COAD KIRC LGG ESCA OV STAD LUAD
110 23 28 31 33 35 36 37 38 38 38 39 40 40 40 41 44 45 50 58
c
BLCA-CESC-HNSC-LUSC
GBM-LGG-PCPG
KIRP-KICH-KIRC
KIRC KICH PCPG LGG GBM THCA STAD ESCA OV LAML ACC TGCT UVM MESO DLBC SARC UCS SKCM BRCA THYM LUSC HNSC CESC BLCA KIRP CHOL PAAD PRAD LIHC LUAD UCEC COAD READ
Pearson's r
-0.4 -0.2 0.0 0.2 0.4 0.6 0.8 1.0
e
ZNF770 DDIT3 SIX4 ZNF135 FOXL1 NR1H2 ZSCAN21 MXI1 ZNF460 CRX ZNF416 HMGA1 ZFP14 HIC1 HMBOX1 IRF1 PRDM9 KLF7 ARID3B KLF4 RUNX3 ZNF213 TBX5 TFAP2A ZNF354A TCF4 ZFX SP1 E2F6 PRDM5 YY1 ZNF354C NFAT5 STAT2 ZBTB24 ZNF263
ACC BLCA BRCA CESC CHOL COAD DLBC ESCA GBM HNSC KICH KIRC KIRP LAML LGG LIHC LUAD LUSC MESO OV PAAD PCPG PRAD READ SARC SKCM STAD TGCT THCA THYM UCEC UCS UVM
Percentage (%)
30 35 40 45 50
f
****
Averged O/E
0.33
0.30
0.27
Real
Random
g
eRNA Number
4
3
2
Gene Number
1
2
3
4
KIRC HNSC GBM ESCA COAD CHOL CESC BRCA UVM UCEC THYM THCA TGCT STAD SKCM SARC READ PRAD PCPG PAAD OV LUSC LUAD LIHC KIRP
i
O/E
0.0 0.5 1.0 1.5
ADORA2A
ADORA2Ae
h
O/E
0.0 1.0 2.0 3.0
GNA11
GNA11e
O/E
0.00 0.25 0.50
ERBB2e
ERBB2
j
Myc
Cell cycle
RTK-RAS
PI3K
Nrf2
IC
Notch
Tp53
Hippo
TGF-beta
CAG
Wnt
eRNA Number
60
40
20
Gene Number
20
40
60

**Extended Data Fig. S4.** eRNAformer discovers novel eRNAs that holds potential value for cancer therapy, related to **Fig. 4**.

**a)** Radar plot displaying the number of detectable eRNAs and their average expression level across different cancer types.

**b)** Pie chart illustrating the distribution of eRNAs based on the number of cancer types in which they are expressed: 1 cancer type, 2–5 cancer types, 6–10 cancer types, and >10 cancer types.

**c)** Heatmap showing the similarity of eRNA expression across tumor samples. A color bar indicates the cancer types, and a scale bar denotes the degree of expression similarity among TCGA samples.

**d)** Radar plot presenting the top 20 cancer types with the highest numbers of master transcription factors (TFs), along with the percentage these TFs represent out of all TFs regulating eRNA expression in the respective cancer type.

**e)** Heatmap of general master regulators across human cancers. The x-axis shows the symbols of master regulators, and the y-axis represents cancer types. General master regulators associated with genomic instability are highlighted in red. The scale bar indicates the percentage of eRNAs correlated with each TF (x-axis) across cancer types (y-axis).

**f)** Bar plot comparing the average Hi-C observed/expected (O/E) contact frequencies between all true eRNA–target links and distance-matched random pairs. *P*-value < 0.001(***), two-sided Wilcoxon signed-rank test. Data are presented as mean ± SEM.

**g)** Radar plot showing the number of eRNAs and immune checkpoint (IC) genes involved in Hi-C-supported regulatory links across various cancer types.

**h)** Hi-C O/E contact frequencies for two regulatory links between eRNAs and clinically actionable genes.

**i)** Hi-C O/E contact frequencies for one regulatory link between an eRNA and an immune checkpoint gene.

**j)** Radar plot indicating the number of genes significantly associated with GDSC anticancer drug sensitivity across different gene sets, along with the number of linked eRNAs.

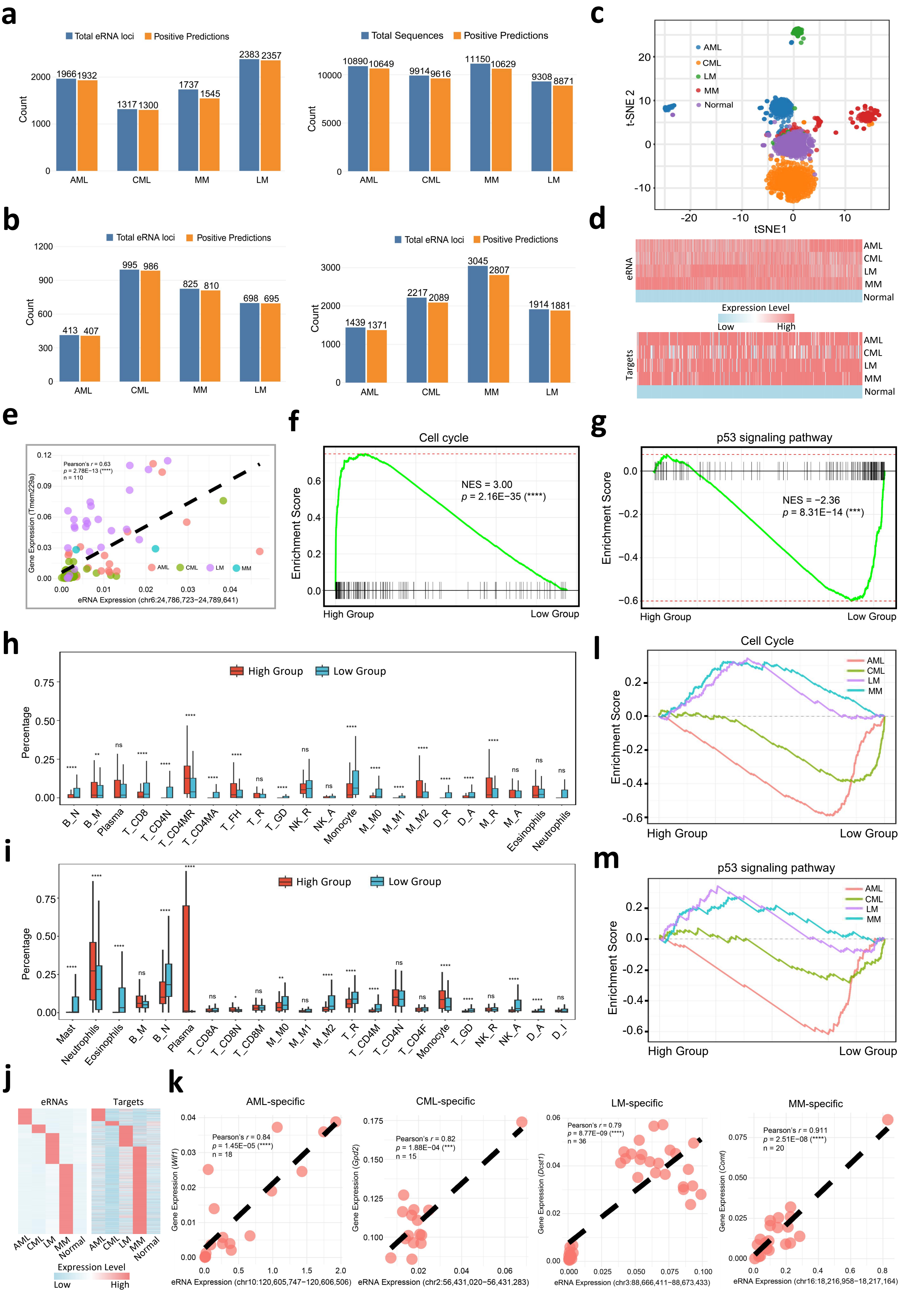
a
Total eRNA loci
Positive Predictions
1966 1932
1317 1300
1737
1545
2383 2357
Count
AML
CML
MM
LM
Total Sequences
Positive Predictions
10890 10649
9914 9616
11150 10629
9308 8871
b
Total eRNA loci
Positive Predictions
413 407
995 986
825 810
698 695
1439 1371
2217 2089
3045 2807
1914 1881
c
AML
CML
LM
MM
Normal
t-SNE 2
tSNE1
d
eRNA
Expression Level
Low
High
Targets
Normal
e
Pearson's r = 0.63
p = 2.78E−13 (****)
n = 110
Gene Expression (Tmem229a)
eRNA Expression (chr6:24,786,723−24,789,641)
f
Cell cycle
NES = 3.00
p = 2.16E−35 (****)
Enrichment Score
High Group
Low Group
g
p53 signaling pathway
NES = −2.36
p = 8.31E−14 (***)
h
High Group
Low Group
Percentage
B_N
B_M
Plasma
T_CD8
T_CD4N
T_CD4MR
T_CD4MA
T_FH
T_R
T_GD
NK_R
NK_A
Monocyte
M_M0
M_M1
M_M2
D_R
D_A
M_R
M_A
Eosinophils
Neutrophils
l
Cell Cycle
i
Mast
Neutrophils
Eosinophils
B_M
B_N
Plasma
T_CD8A
T_CD8N
T_CD8M
M_M0
M_M1
M_M2
T_R
T_CD4M
T_CD4N
T_CD4F
Monocyte
T_GD
NK_R
NK_A
D_A
D_I
m
p53 signaling pathway
j
eRNAs
Targets
Expression Level
k
AML-specific
Pearson's r = 0.84
p = 1.45E−05 (****)
n = 18
Gene Expression (Wif1)
eRNA Expression (chr10:120,605,747−120,606,506)
CML-specific
Pearson's r = 0.82
p = 1.88E−04 (***)
n = 15
Gene Expression (Gpd2)
eRNA Expression (chr2:56,431,020−56,431,283)
LM-specific
Pearson's r = 0.79
p = 8.77E−09 (****)
n = 36
Gene Expression (Dcst1)
eRNA Expression (chr3:88,666,411−88,673,433)
MM-specific
Pearson's r = 0.911
p = 2.51E−08 (****)
n = 20
Gene Expression (Comt)
eRNA Expression (chr16:18,216,958−18,217,164)

**Extended Data Fig. S5.** A multitude of cancer-related eRNAs are aberrantly activated in hematological malignancies, related to **Fig. 5**.

**a)** Barplot showing the number of mono-exon transcript loci that overlap with known eRNAs in different human hematological malignancies, alongside the number of these loci predicted as authentic eRNAs. Left, known eRNAs from FANTOM5; Right, known eRNAs from eRNAbase.

**b)** Barplot showing the number of mono-exon transcript loci that overlap with known eRNAs in different mouse hematological malignancies, alongside the number of these loci predicted as authentic eRNAs. Left, known eRNAs from FANTOM5; Right, known eRNAs from eRNAbase.

**c)** t-SNE plot illustrating the projection of mouse hematologic tumor samples and normal samples onto the first two dimensions, with colors indicating sample types.

**d)** Heatmap displaying 1,525 consistently activated eRNAs and their 431 target genes across multiple mouse hematological malignancies.

**e)** Scatter plot showing the expression correlation between a consistently activated eRNA (chr6:24,786,723-24,789,641) and the cancer-related gene *Tmem229a* across different mouse hematological malignancies. The *p*-value was obtained from a correlation coefficient test.

**f)** GSEA plot showing the expression changes of cell cycle genes between high-score and low-score groups stratified by consistently activated eRNA target genes in human hematological malignancies.

**g)** GSEA plot showing the expression changes of p53 signaling pathway genes between high-score and low-score groups stratified by consistently activated eRNA target genes in mouse hematological malignancies.

**h)** Violin plot showing the abundance distribution of different cell types between high-score and low-score groups stratified by consistently activated eRNA target genes in human hematological malignancies. B_N, Naive B-cells; B_M, Memory B-cells; T_CD8, CD8 T-cells; T_CD4N, CD4 Naive T-cells; T_CD4MR, CD4 Memory Resting T-cells; T_CD4MA, CD4 Memory Activated T-cells; T_FH, Follicular Helper T-cells; T_R, Regulatory T-cells; T_GD, Gamma Delta T-cells; NK_R, Resting NK cells; NK_A, Activated NK cells; M_M0, M0 Macrophages; M_M1, M1 Macrophages; M_M2, M2 Macrophages; D_R, Resting Dendritic cells; D_A, Activated Dendritic cells; M_R, Resting Mast cells; M_A, Activated Mast cells.

**i)** Violin plot showing the abundance distribution of different cell types between high-score and low-score groups stratified by consistently activated eRNA target genes in mouse hematological malignancies.B_N, Naive B-cells; B_M, Memory B-cells; T_CD8A, CD8 Activated T-cells; T_CD8N, CD8 Naive T-cells; T_CD8M, CD8 Memory T-cells; M_M0, M0 Macrophages; M_M1, M1 Macrophages; M_M2, M2 Macrophages; T_R, Regulatory T-cells; T_CD4M, CD4 Memory T-cells;T_CD8N, CD8 Naive T-cells; T_CD4F, CD4 Follicular T-cells; T_GD, Gamma Delta T-cells; D_R, Resting Dendritic cells; D_A: Activated Dendritic cells; D_I, Immature Dendritic cells.

**j)** Heatmap displaying 10,037 eRNAs (AML: 1,283; CML: 687; LM: 2,466; MM: 5,601) and 2,158 target genes (AML: 217; CML: 77; LM: 361; MM: 1,503) specifically expressed in distinct mouse hematological malignancies.

**k)** Scatter plot showing the expression correlations between four cancer-type-specific eRNAs and their target genes in mouse hematological malignancies. The *p*-value was derived from a correlation coefficient test.

**l)** GSEA plot showing the expression changes of cell cycle genes between high-score and low-score groups stratified by cancer-specific expressed eRNA target genes in mouse hematological malignancies.

**m)** GSEA plot showing the expression changes of p53 signaling pathway genes between high-score and low-score groups stratified by cancer-specific expressed eRNA target genes in mouse hematological malignancies.

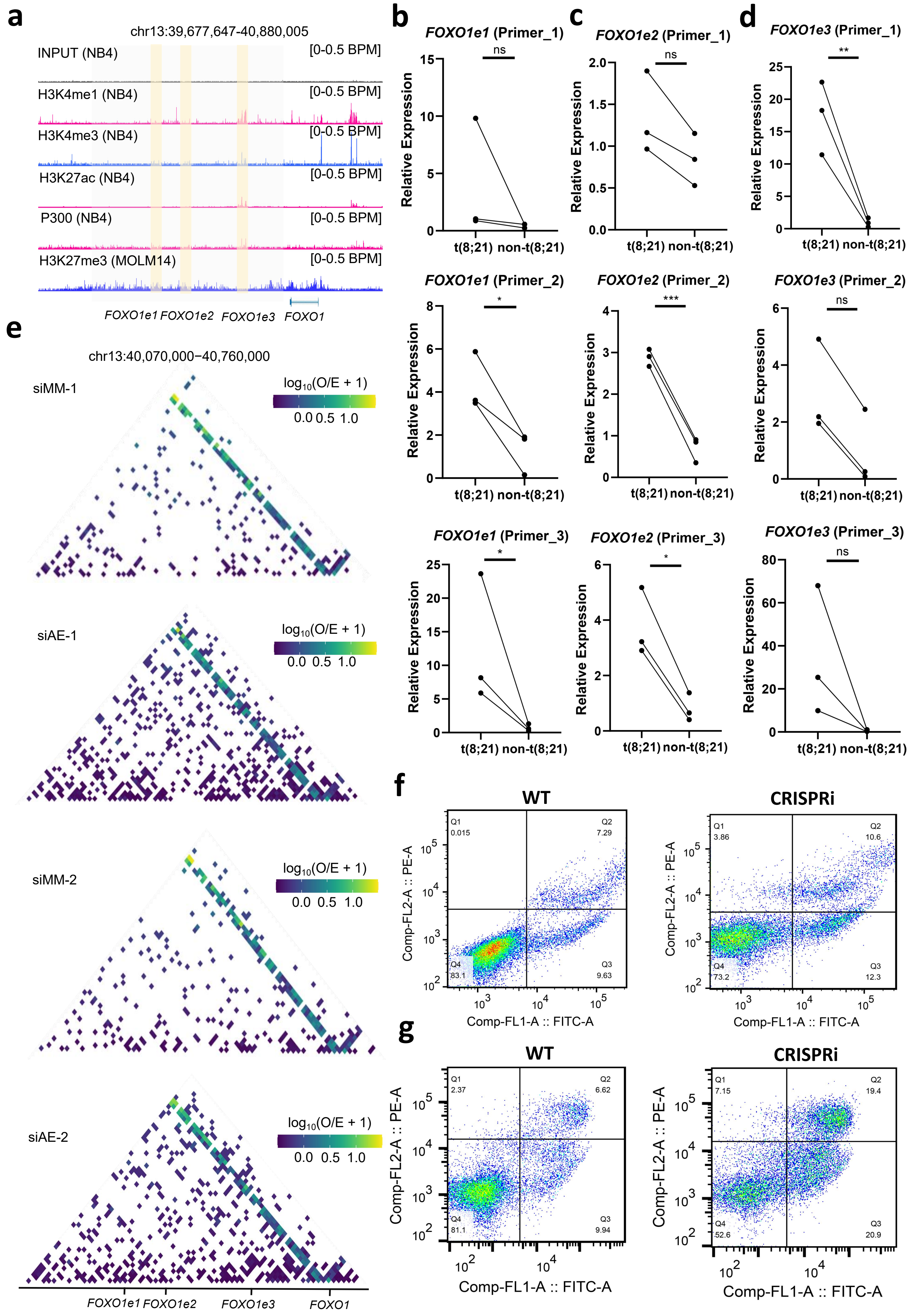
a
chr13:39,677,647-40,880,005
INPUT (NB4)
[0-0.5 BPM]
H3K4me1 (NB4)
H3K4me3 (NB4)
H3K27ac (NB4)
P300 (NB4)
H3K27me3 (MOLM14)
FOXO1e1 FOXO1e2 FOXO1e3 FOXO1
b
FOXO1e1 (Primer_1)
FOXO1e1 (Primer_2)
FOXO1e1 (Primer_3)
c
FOXO1e2 (Primer_1)
FOXO1e2 (Primer_2)
FOXO1e2 (Primer_3)
d
FOXO1e3 (Primer_1)
FOXO1e3 (Primer_2)
FOXO1e3 (Primer_3)
Relative Expression
t(8;21) non-t(8;21)
ns
*
**
***
e
chr13:40,070,000−40,760,000
siMM-1
siAE-1
siMM-2
siAE-2
log10(O/E + 1)
0.0 0.5 1.0
FOXO1e1 FOXO1e2 FOXO1e3 FOXO1
f
g
WT
CRISPRi
Comp-FL2-A :: PE-A
Comp-FL1-A :: FITC-A
Q1 0.015
Q2 7.29
Q3 9.63
Q4 83.1
Q1 3.86
Q2 10.6
Q3 12.3
Q4 73.2
Q1 2.37
Q2 6.62
Q3 9.94
Q4 81.1
Q1 7.15
Q2 19.4
Q3 20.9
Q4 52.6

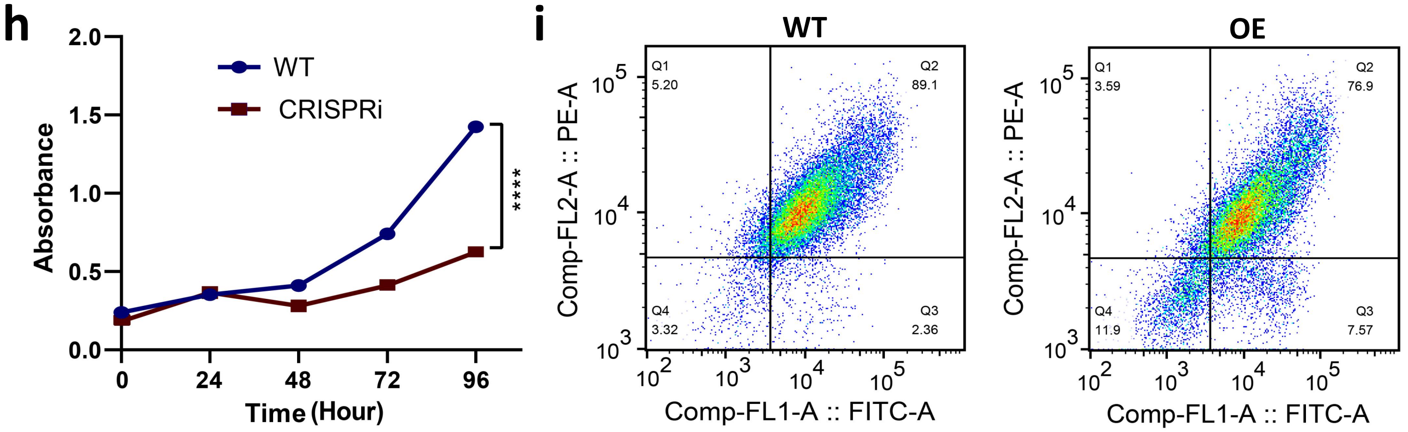


**Extended Data Fig. S6.** *FOXO1e* is an eRNA cluster that promotes t(8;21) AML oncogenesis, related to **Fig. 6**.

**a)** ChIP-seq was used to profile the enrichment of H3K4me1, H3K4me3, H3K27ac, P300, and H3K27me3 within and flanking the *FOXO1e* locus. Signal intensities are BPM-normalized and presented as read density tracks.

**b)** Heatmap visualization of alterations in Hi-C interactions following perturbation of AE fusion gene expression.

**c)** *FOXO1e1* expression levels were quantified by qPCR using three independent primer sets in a cohort of t(8;21) AML patients versus non-t(8;21) AML patients. Statistical significance was assessed by two-way ANOVA. $p$-value < 0.05 (*), $p$-value > 0.05 (ns).

**d)** *FOXO1e2* expression levels measured by qPCR with three distinct primer sets in three t(8;21) AML patients and non-t(8;21) AML patients. $p$-value < 0.001 (***), $p$-value < 0.05 (*), $p$-value > 0.05 (ns), two-way ANOVA test.

**e)** *FOXO1e3* expression levels were similarly evaluated by qPCR using three independent primer sets across the same patient groups. Significance determined by two-way ANOVA, $p$-value < 0.01 (**), $p$-value > 0.05 (ns), two-way ANOVA test.

**f)** Flow cytometric analysis revealed a pronounced increase in apoptotic cell fractions in *FOXO1e3*-knockdown cells relative to control cells at 72 hours post-transfection.

g) *FOXO1e3*-knockdown cells exhibited a more obvious increase in apoptosis levels compared to control cells at 96 hours post-transfection, as determined by flow cytometry.

**h)** A dot-line plot demonstrates a significant decrease in proliferation of *FOXO1e3*-knockdown cells relative to controls.

**i)** Flow cytometry analysis at 72 hours post-transfection showed that *FOXO1e3*-overexpressing cells undergo a substantial reduction in apoptosis compared to control cells.
